# Multi-genome comparisons reveal gain-and-loss evolution of the *anti-Mullerian hormone receptor type 2* gene, an old master sex determining gene, in Percidae

**DOI:** 10.1101/2023.11.13.566804

**Authors:** Heiner Kuhl, Peter T Euclide, Christophe Klopp, Cedric Cabau, Margot Zahm, Céline Roques, Carole Iampietro, Claire Kuchly, Cécile Donnadieu, Romain Feron, Hugues Parrinello, Charles Poncet, Lydia Jaffrelo, Carole Confolent, Ming Wen, Amaury Herpin, Elodie Jouanno, Anastasia Bestin, Pierrick Haffray, Romain Morvezen, Taina Rocha de Almeida, Thomas Lecocq, Bérénice Schaerlinger, Dominique Chardard, Daniel Żarski, Wes Larson, John H. Postlethwait, Serik Timirkhanov, Werner Kloas, Sven Wuertz, Matthias Stöck, Yann Guiguen

**Author notes:** corresponding authors: Heiner Kuhl and Yann Guiguen.

## Abstract

The Percidae family comprises many fish species of major importance for aquaculture and fisheries. Based on three new chromosome-scale assemblies in *Perca fluviatilis*, *Perca schrenkii* and *Sander vitreus* along with additional percid fish reference genomes, we provide an evolutionary and comparative genomic analysis of their sex-determination systems. We explored the fate of a duplicated anti-Mullerian hormone receptor type-2 gene (*amhr2bY*), previously suggested to be the master sex determining (MSD) gene in *P. flavescens*. Phylogenetically related and structurally similar a*mhr2* duplications (*amhr2b*) were found in *P. schrenkii* and *Sander lucioperca*, potentially dating this duplication event to their last common ancestor around 19-27 Mya. In *P. fluviatilis* and *S. vitreus*, this *amhr2b* duplicate has been lost while it was subject to amplification in *S. lucioperca*. Analyses of the *amhr2b* locus in *P. schrenkii* suggest that this duplication could be also male-specific as it is in *P. flavescens*. In *P. fluviatilis*, a relatively small (100 kb) non-recombinant sex-determining region (SDR) was characterized on chromosome-18 using population-genomics approaches. This SDR is characterized by many male-specific single-nucleotide variants (SNVs) and no large duplication/insertion event, suggesting that *P. fluviatilis* has a male heterogametic sex determination system (XX/XY), generated by allelic diversification. This SDR contains six annotated genes, including three (*c18h1orf198*, *hsdl1*, *tbc1d32*) with higher expression in testis than ovary. Together, our results provide a new example of the highly dynamic sex chromosome turnover in teleosts and provide new genomic resources for Percidae, including sex-genotyping tools for all three known *Perca* species.

## INTRODUCTION

The percid family (Percidae, Rafinesque) encompasses a large number (over 250) of diverse ecologically and economically important fish species, assigned to 11 genera [1]. Two genera, *Perca* and *Sander* are found across both Eurasia and North America, with separate species native to each continent (Eurasia: *Perca fluviatilis / Sander lucioperca*; North America: *Perca flavescens / Sander vitreus*). Percids are classically described as typical freshwater species of the Northern hemisphere, even if some species can be regularly found in brackish waters (e.g. *Sander lucioperca, Perca fluviatilis*). In the context of declining fisheries over the past few decades, but also due to their high value and good market acceptance, four percid species - *Perca flavescens* (yellow perch) and *Sander vitreus* (walleye) in North America and *P. fluviatilis* (European perch) and *S. lucioperca* (zander) in Eurasia are particularly promising for aquaculture. Rearing these fish in recirculation aquaculture systems (RAS) allows for a control of reproduction and a year-round production of stocking fish [2,3]. Although year-round production represents an important competitive goal, current production targets premium markets and an up-scaling of production faces several bottlenecks [4].

Among these bottlenecks is better control of the sex of developing individuals because in both *Perca* and *Sander* genera, females grow faster than males [5,6]. Due to faster female growth (up to 25-50% in *Perca*, 10% in *Sander*), all-female stocks are highly desirable. In *Perca fluviatilis*, sex determination has been assumed to be male heterogametic (XX/XY) based on gynogenesis or hormonal treatment experiments [7,8]. These methodologies also produced genetic female but phenotypic male individuals (neomales) that can be used to produce all-female stocks by crossing normal XX females with these chromosomally XX neomales. This approach would, however, greatly benefit from a reliable sexing method allowing the identification of genetic sex early during development to select rare genetically XX neomales as future breeders in aquaculture. In *P. flavescens,* an XX female/XY male heterogametic genetic sex determination system has been also recently uncovered, with duplication / insertion of an anti-Mullerian hormone receptor type 2 (*amhr2*) gene as a potential master sex determining gene [9].

Genes encoding many members of the transforming growth factor beta (TGF-β) gene family, including anti-Mullerian hormone (*amh*) and anti-Mullerian hormone receptor type-2 (*amhr2*), have repeatedly and independently evolved as master sex-determining (MSD) genes in vertebrates [10]. For instance, *amh* has been characterized or suspected to be the MSD gene in pikes [11,12], Nile tilapia [13], lumpfish [14], *Sebastes* rockfish [15], lingcod [16], and Patagonian pejerrey [17]. The cognate receptor gene of Amh, *amhr2,* has also been found as a potential MSD gene in Pangasiidae [18], Takifugu [19], Ayu [20], common seadragon and alligator pipefish [21], as well as in yellow perch [9]. The repeated and independent recruitment of TGF-β receptors, including *Amhr2*, in teleost fish sex determination is even more puzzling as many of these MSD genes, encoding a TGFβ receptor, share a similar N-terminal truncation [9,18,21], supporting their evolution towards a ligand-independent mechanism of action [18]. Therefore, the extent of evolutionary conservation of the Y-linked *amhr2bY* gene found in yellow perch in closely related species (genus *Perca* and *Sander*) is an important question with implications for better sex-control in these aquaculture species and for understanding the evolution of sex linkage and protein structure.

Regarding genomics of Percidae, two long-read reference quality genome assemblies have recently been published for *P. flavescens* and *S. lucioperca* [9,22]. While for *P. fluviatilis* and *S. vitreus* only draft genomes, generated from short-read sequencing, have been available [23]. Here, we provide three new long-read chromosome-scale genome assemblies for *P. fluviatilis*, *P. schrenkii* and *Sander vitreus* and thus complete genomic resources for the economically most important species of Percidae. These data enabled us to develop PCR-assays for sexing of all three *Perca* species and shed light on gene gain-and-loss in the evolution of an old MSD gene in Percidae.

## MATERIAL AND METHODS

### Biological samples

In *Perca fluviatilis,* high molecular weight (HMW) genomic DNA (gDNA) for genome sequencing was extracted from a blood sample of a male called “Pf_M1” (BioSample ID SAMN12071746) from the aquaculture facility of the University de Lorraine, Nancy, France. Blood (0.5 ml) was sampled and directly stored in 25 ml of a TNES-Urea lysis buffer (TNES-Urea: 4 M urea; 10 mM Tris-HCl, pH 7.5; 125 mM NaCl; 10 mM EDTA; 1% SDS). HMW gDNA was extracted from the TNES-urea buffer using a slightly modified phenol/chloroform protocol as described [12]. For the chromosome contact map (Hi-C), 1.5 ml of blood was taken from the same animal and slowly (1 K/min) cryopreserved with 15 % dimethyl sulfoxide (DMSO) in a Mr. Frosty Freezing Container (ThermoFisher) at −80°C. Additional fin clip samples for RAD-Sequencing (RAD-Seq), Pool-Sequencing (Pool-Seq) or sex-genotyping assays were collected and stored in 90% ethanol, either at the Lucas Perche aquaculture facility (Le Moulin de Cany, 57170 Hampont, France), at Kortowskie Lake in Poland, or at Mueggelsee Lake in Germany.

Samples of *Perca schrenkii* were obtained for genome sequencing and sex genotyping from male and female wild catches at lake Alakol, Kazakhstan (46.328 N, 81.374 E). Different organs and tissues (brain, liver, muscle, ovary, testis) were sampled for genome and transcriptome sequencing (Biosample ID SAMN15143703) and stored in RNAlater. HMW gDNA for genome sequencing was extracted from brain tissue of the male *P. schrenkii* individual, using the MagAttract HMW DNA Kit (Qiagen, Germany). Total RNA for transcriptome sequencing was isolated using a standard Trizol protocol, in combination with the RNAeasy Mini Kit (Qiagen, Germany).

For genome sequencing of *Sander vitreus* a fin clip of a male was sampled by Ohio Department of Natural Resources (Ohio, DNR) in spring 2017 and stored in 96% ethanol. The *S. vitreus* sample called “19-12246” originated from Maumee River, Ohio [41.554 N; −83.6605W]. DNA was extracted using the DNeasy Tissue Kit (Qiagen). Short DNA fragments were removed/reduced by size-selective, magnetic-bead purification using 0.35x of sample volume AMPure beads (Beckmann-Coulter) and two washing steps with 70% ethanol.

### Sequencing

Genomic sequencing of *P. fluviatilis* was carried out using a combination of 2x250 bp Illumina short-reads, Oxford Nanopore long reads and a chromosome contact map (Hi-C). For long-read sequencing, DNA was sheared to 20 kb using the megaruptor system (Diagenode). ONT (Oxford nanopore technologies) library preparation and sequencing was performed using 5 µg of sheared DNA and ligation sequencing kits SQK-LSK108 or SQK-LSK109, according to the manufacturer’s instructions. The libraries were loaded at a concentration of 0.005 to 0.1 pmol and sequenced for 48 h on 11 GridION R9.4 or R9.4.1 flowcells. Short read wgs (whole genome shotgun) sequencing for consensus polishing of noisy long read assemblies was carried out by shearing the HMW DNA to approximately 500 bp fragments and using the Illumina Truseq X kit, according to the manufacturer’s instructions. The library was sequenced using a read length of 250 bp in paired-end mode (HiSeq 3000, Illumina, California, USA). Hi-C library generation for chromosome assembly was carried out according to a protocol adapted from Rao *et al*. 2014 [24]. The blood sample was spun down, and the cell pellet was resuspended and fixed in 1% formaldehyde. Five million cells were processed for the Hi-C library. After overnight-digestion with *Hind*III (NEB), DNA-ends were labeled with Biotin-14-DCTP (Invitrogen), using Klenow fragment (NEB) and re-ligated. A total of 1.4 µg of DNA was sheared to an average size of 550 bp (Covaris). Biotinylated DNA-fragments were pulled down using M280 Streptavidin Dynabeads (Invitrogen) and ligated to PE adaptors (Illumina). The Hi-C library was amplified using PE primers (Illumina) with 10 PCR amplification cycles. The library was sequenced using a HiSeq3000 (Illumina, California, USA), generating 150 bp paired-end reads.

Genomic sequencing of *P. schrenkii* and *S. vitreus* was carried out using Oxford Nanopore long reads on a MinION nanopore sequencer (Oxford Nanopore Technologies, UK) in combination with the MinIT system. Several libraries were constructed using the tagmentation-based SQK-RAD004 kit with varying amounts of input DNA (0.4 to 1.2 µg) from a male individual or using the ligation approach of the SQK-LSK109 kit (input DNA 2 µg). Libraries were sequenced on R9.4.1 flowcells with variable run times and exonuclease washes by the EXP-WSH003 kit to remove pore blocks and improve the data yield. Short-read wgs-sequencing of *P. schrenkii* was conducted at BGI (BGI Genomics Co., Ltd.). A *P. schrenkii* male and a female wgs library (300 bp fragment length) were constructed and paired end reads of 150 bp length were generated on an Illumina Hiseq4000 system. Public short-read wgs data of *S. vitreus* were obtained from the NCBI Sequence Read Archive (SRA) using the accession SRR9711286. Transcriptome sequencing of six *P. schrenkii* samples (female brain, male brain, male muscle, female liver, ovary and testis) was conducted at BGI. Transcriptome-sequencing libraries were constructed from total RNA, applying enrichment of mRNA with oligo(dT) hybridization, mRNA fragmentation, random hexamer cDNA synthesis, size selection and PCR amplification. Sequencing of 150 bp paired-end reads was performed by an Illumina HiSeq X Ten system.

### Genome assembly of Perca fluviatilis

Residual adaptor sequences in ONT GridION long reads were trimmed and split by Porechop (v0.2.1) [25]. Reads longer than 9999 bp were assembled by SmartDeNovo (May-2017) [26] using default parameters. Long reads were remapped to the SmartDeNovo contigs by Minimap2 (v2.7) [27] and Racon (v1.3.12) [28] was used to polish the consensus sequence. In a second round of polishing, Illumina short-reads were mapped by BWA mem (v0.7.12-r1039) [29] to the contigs, which were subsequently polished by Pilon (v1.223) [30]. The chromosome-scale assembly was performed by mapping Hi-C data to the assembled contigs, using the Juicer pipeline (v1.5.6) [31] and subsequent scaffolding by 3D-DNA (v180114) [32]. Juicebox (v1.8.8) [33] was used to manually review and curate the chromosome-level scaffolds. A final gap-closing step, applying long reads and LR_gapcloser (v1.1, default parameters) [34], further increased contig length. After gap-closing, a final consensus sequence polishing step was performed by mapping short reads to the scaffolds, sequence variants (1/1 genotypes were considered as corrected errors) were detected with Freebayes (v0.9.7) [35] and written to a vcf-file. The final fasta file was then generated by vcf-consensus from Vcftools (v0.1.15, default parameters).

### Genome assembly of Perca schrenkii *and* Sander vitreus

Illumina short reads were trimmed using Trimmomatic (v0.35) [36]. Short reads were assembled using a custom compiled high kmer version of idba-ud (v1.1.1) [37] with kmer size up to 252. The resulting contigs were mapped against available Percidae genomes (*P. flavescens, P. fluviatilis* and *S. lucioperca*) by Minimap2 and analysis of overall mapped sequence length resulted in *P. schrenkii* aligned best with *P. flavescens* and *S. vitreus* aligned best with *S. lucioperca*. According to the benchmarks published in [38], the publicly available chromosome-level assembly of *P. flavescens* (RefSeq: GCF_004354835.1) could be used to aid the chromosome assembly of *P. schrenkii* as follows: ONT MinION long reads (male sample) were trimmed and split using Porechop (v0.2.1) [25]. The inhouse developed CSA method (v2.6) [38], was used to assemble the *P. schrenkii* genome from long-read data and short-read contigs and to infer chromosomal scaffolds using the *P. flavescens* reference genome. CSA parameters were optimized to account for relatively low long-read sequencing coverage and hybrid assembly of long reads and short-read contigs:

CSA2.6.pl –r longreads.fa.gz –g P.flavescens.fa –k 19 –s 2 –e 2 –l „–i shortreadcontigs.fa –L3000 –A“

Similarly, we assembled *S. vitreus*, using the *S. lucioperca* contigs and *P. flavescens* chromosomes as references for chromosomal assembly. Here, the short-read contigs were treated as long-reads:

CSA2.6c.pl -r longreads+contigs.fa.gz -g sanLuc.CTG.fa.gz,PFLA_1.0_genomic.fna.gz -k 19 -s 2 -e 2

The assemblies were manually curated, and the consensus sequences were polished using long reads and flye (v2.6) [39], with options: --nanoraw --polish-target, followed by two rounds of polishing by Pilon (v1.23) [30], using the short-read data, which had been mapped by Minimap2 (v2.17-r941) [27], to the genome assemblies.

### Genome annotation

*De novo* repeat annotation was performed using RepeatModeler (version open-1.0.8) and Repeat Masker (version open-4.0.7). The *P. fluviatilis* genome has been assigned to the RefSeq assembly section of NCBI and has been annotated by GNOMON (www.ncbi.nlm.nih.gov/genome/annotation_euk/process), which included evidence from Actinopterygii proteins (n=154,659) and *P. fluviatilis* RNAseq reads (n = 3,537,868,978) (www.ncbi.nlm.nih.gov/genome/annotation_euk/Perca_fluviatilis/100). To annotate our *P. schrenkii* and *S. vitreus* assemblies, we used the high-quality GNOMON annotations from their closest relatives *P. flavescens* (www.ncbi.nlm.nih.gov/genome/annotation_euk/Perca_flavescens/100) and *S. lucioperca* (www.ncbi.nlm.nih.gov/genome/annotation_euk/Sander_lucioperca/101), respectively. We performed high-throughput comparative protein coding gene annotation by spliced alignment of GNOMON mRNAs and proteins by Spaln (v2.06f, [40]) to our assemblies and combined the resulting CDS- and UTR-matches into complete gene models by custom scripts. All annotations were benchmarked using BUSCO [41] with the Actinopterygii_odb9 database and obtained highly similar values as the reference annotations used for the comparative annotation approach.

### Genome browsers and data availability

We provide UCSC genome browsers [42] for the five available *Perca* and *Sander* reference genomes (this study: *P. fluviatilis*, *P. schrenkii*, *S. vitreus*; earlier studies: *P. flavescens* [9] and *S. lucioperca* [22] at http://genomes.igb-berlin.de/Percidae/. These genome browsers provide access to genomic sequences and annotations (either public NCBI GNOMON annotations or annotations resulting from our comparative approach). Blat [43] servers for each genome are available to align nucleotide or protein sequences.

### Phylogenomics and divergence time estimation

We performed pair-wise whole-genome alignments of 36 teleost genome assemblies as in [44], using Last-aligner and Last-split [45] for filtering 1-to-1 genome matches, Multiz [46] for multiple alignment construction from pairwise alignments and filtered for non-coding sequences to calculate the species tree using iqtree2 and raxml-ng [47,48]. We added the genomes *of P. schrenkii, S. vitreus and Etheostoma spectabile* (GCF_008692095.1) to this dataset and re-analyzed the highly-supported subclade containing Percidae species using several outgroups (*Lates, Oreochromis, Pampus* and *Thunnus sp.).* We estimated divergence times using a large subset of our multiple alignment (10^6^ nt residues) and the approximate method of Mcmctree (Paml package version, [49]). We calibrated 5 nodes of the tree by left or right CI values, obtained from www.timetree.org and applied independent rates or correlated rates clock models and the HKY85 evolutionary model. Approximately 10^8^ samples were calculated, of which we used the top 50% for divergence time estimation. Each calculation was performed in two replicates, which were checked for convergence using linear regression. The final tree was plotted using FigTree (v1.4.4, http://tree.bio.ed.ac.uk/software/figtree).

### Perca fluviatilis RAD-Sequencing

*Perca fluviatilis* gDNA samples from 35 males and 35 females were extracted with the NucleoSpin Kit for Tissue (Macherey-Nagel, Duren, Germany), following the manufacturer’s instructions. Then, gDNA concentrations were quantified with a Qubit3 fluorometer (Invitrogen, Carlsbad, CA) using a Qubit dsDNA HS Assay Kit (Invitrogen, Carlsbad, CA). RAD libraries were constructed from each individual’s gDNA, using a previously described protocol with the single *Sbf*1 restriction enzyme [50]. These libraries were sequenced on an Illumina HiSeq 2500. Raw reads were demultiplexed using the process_radtags.pl wrapper script of stacks, version 1.44, with default settings [51], and further analyzed with the RADSex analysis pipeline [52] to identify sex-specific markers.

### Perca fluviatilis Pool-Sequencing

Sequencing of pooled samples (Pool-Seq) was carried out in *Perca fluviatilis* to increase the resolution of RAD-Sequencing for the identification of sex-specific signatures characteristic of its sex-determining region. The gDNA samples used for RAD-Sequencing were pooled in equimolar quantities according to their sex. Pooled male and pooled female libraries were constructed using a Truseq nano kit (Illumina, ref. FC-121-4001) following the manufacturer’s instructions. Each library was sequenced in an Illumina HiSeq2500 with 2x 250 reads. Pool-Seq reads were analyzed as previously described [9,11,53–55] with the PSASS pipeline (psass version 2.0.0: https://zenodo.org/record/2615936#.XtyIS3s6_AI) that computes the position and density of single nucleotide variations (SNVs), heterozygous in one sex but homozygous in the other sex (sex-specific SNVs), and the read depths for the male and female pools along the genome to look for sex coverage differences. Psass was run with default parameters except –window-size, which was set to 5,000, and –output-resolution, which was set to 1,000.

### PCR-based sex diagnostics

A *Perca schrenkii* PCR-based sex-diagnostic test was designed based on multiple alignments of the different *amhr2* genes in *P. fluviatilis* (one autosomal gene only)*, P. flavescens* (two genes), and *Perca schrenkii* (two genes) to target a conserved region for all *Perca amhr2* genes, allowing the design of PCR-primers that amplify both the autosomal *amhr2a* and the male-specific *amhr2bY* with different and specific PCR-amplicon sizes. Selected PCR primer sequences were forward: 5’-AGTTTATTGTGTTAGTTTGGGCT-3’ and reverse: 5’-CAAATAAATCAGAGCAGCGCATC-3’. PCRs were carried out with 1U Platinum Taq DNA Polymerase and its corresponding Buffer (Thermofisher) supplemented with 0.8 mM dNTPs (0.2mM each), 1.5 mM MgCl_2_ and 0.2 µM of each primer with the following cycling conditions, 96°C for 3 min; 40 cycles of denaturation (96°C, 30 s), annealing (54°C, 30 s) and extension (72°C, 1 min); final extension (72°C, 5 min); storage at 4°C. PCR amplicons were separated on 1.5% agarose gels (1.5% std. agarose, 1x TBE buffer, 5 V/cm, running time 40 min) and the systematic amplification of the autosomal (*amhr2a*) amplicon was used as a positive PCR control.

*Perca fluviatilis* primers were designed to amplify a 27 bp-deletion variant in the third intron of the *P. fluviatilis hsdl1* gene, which was identified as a male specific (Y-specific) variation based on the pool-seq analysis. Selected PCR-primer sequences were forward 5’-ACACTGATCAACATTTTCTGTCTCA-3’ and reverse 5’-TGTTAACATTTGAGAATTTTGCCTT-3’. PCRs were carried out as described above with the following cycling conditions: denaturation 96°C for 3 min; 40 cycles of denaturation (96°C, 30 s), annealing (60°C, 30 s) and extension (72°C, 30 min); final extension (72°C, 5 min); storage at 4°C. PCR amplicons were separated on 5% agarose gels (5% Biozym sieve 3:1 agarose, 1x TBE buffer, 5 V/cm, 1 h 40 min running time) and the amplicon derived from the amplification of the X-chromosome allele was used as a positive PCR control. In addition to this classical PCR sex-genotyping method, we also explored the sex-linkage of some sex-specific SNVs in *P. fluviatilis* using Kompetitive Allele-Specific Polymerase chain reaction (KASPar) assays [56]. Seven sex-specific SNVs were selected at different locations within the *P. fluviatilis* sex-determining region. Primers (Table 1) were designed using the design service available on the 3CR Bioscience website (www.3crbio.com/free-assay-design). KASPar genotyping assays were carried out with a single end-point measure on a Q-PCR Light Cycler 480 (Roche) using the Agencourt® DNAdvance kit (Beckman), following the manufacturer’s instructions.

**Table 1:**
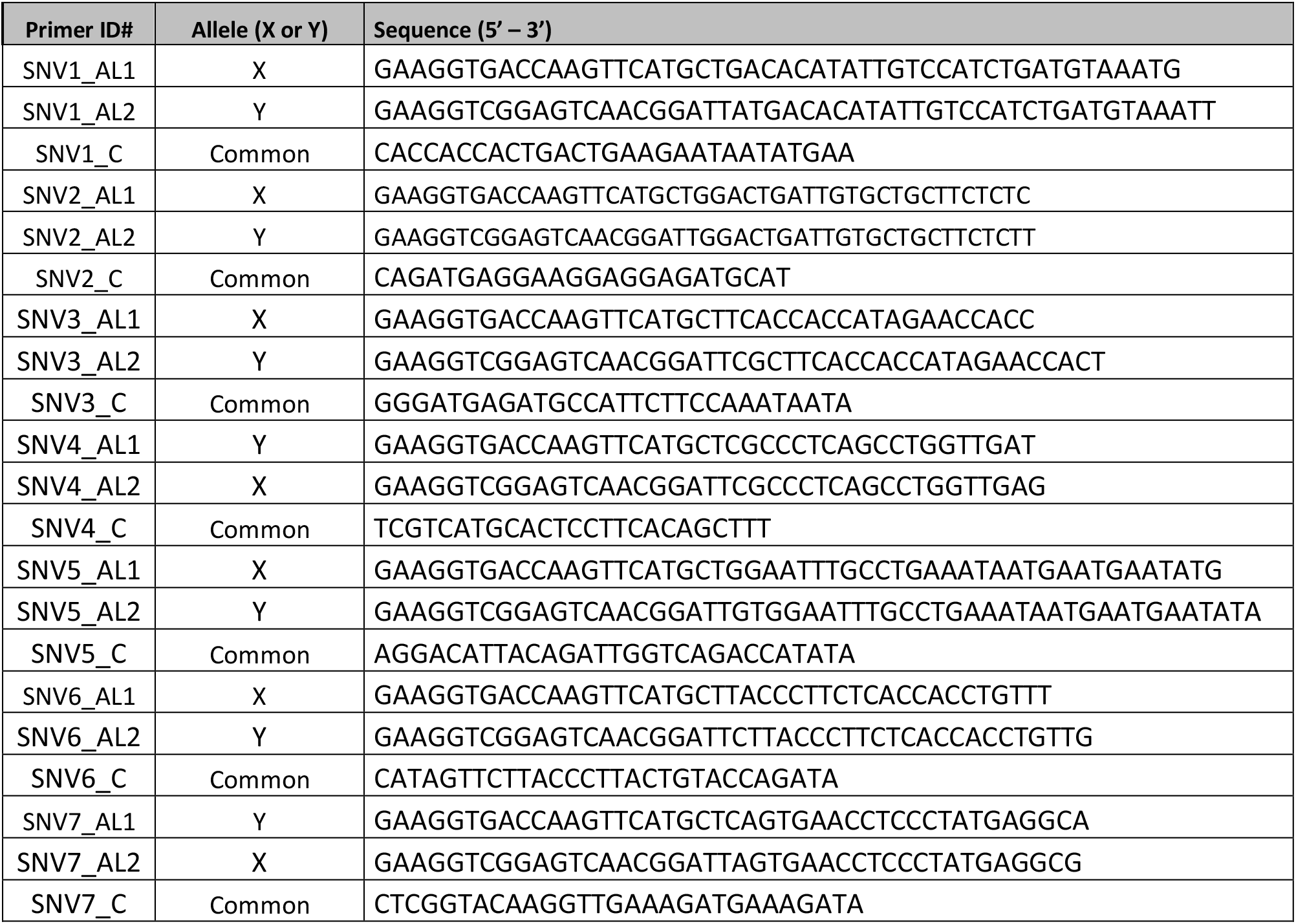
KASpar allele-specific PCR-primers. For each allele (AL1 and AL2) primers, the X and Y sex chromosome-specific alleles are provided along with their sequences.

### RNAseq expression analyses

Already available gonadal datasets of *P. fluviatilis* (age 9 month) RNAseq [57] (SRA accessions: SRR14461526 and SRR14461527) were used to compare gene expression between ovary and testes for the gene models annotated in the SD-region. Reads were mapped to our *P. fluviatilis* reference genome using HISAT2 [58] and transcript assembly and FPKM-values were calculated using STRINGTIE [59].

## RESULTS

### Genome assemblies of Perca fluviatilis, Perca schrenkii and Sander vitreus

The genome of *P. fluviatilis* was sequenced to high coverage using Oxford Nanopore long-read sequencing (estimated coverage: 67-fold / N50 read length: 11.9 kbp), Hi-C data was generated to allow for chromosome-level assembly (coverage: 52-fold / alignable pairs 89.1%/ Hi-C map see Suppl. Fig. 1). The final assembly yielded a highly complete reference genome (99.0% of sequence assigned to 24 chromosomes (N50 length: 39.6 Mbp) and highly continuous contigs (N50 length: 4.1 Mbp). Compared to a previously published genome assembly of *P. fluviatilis,* obtained from “linked-short-reads” (10X Genomics), these numbers represent a 316-fold improvement of contig continuity and a 6.3-fold increase of scaffold continuity. The better continuity resulted in an increased percentage of predictable genes (BUSCO results below). Genome assembly statistics are also highly congruent with the previously published reference quality *P. flavescens* genome, except for obvious size differences as the *P. fluviatilis* assembly is about 8.1% larger than the *P. flavescens* assembly and represents the largest genome known in the genus *Perca* (Table 2).

**Table 2:** Genome assembly statistics for *P. fluviatilis* and *P. schrenkii*. For comparison, the table also provides numbers for two recently published *Perca sp.* assemblies [9,64]. Abbreviations: LR = long reads; HiC = chromosome conformation capture; CG = comparative genomics; 10X = linked-read sequencing; GLM = genetic linkage map.

|  | this study |  |  | earlier studies |  |  |
| --- | --- | --- | --- | --- | --- | --- |
| Species | *P. fluviatilis* | *P. schrenkii* | *S. vitreus* | *P. flavescens* | *P. fluviatilis* | *S. lucioperca* |
| Strategy | LR+HiC | LR+CG | LR+CG | LR+10X+HiC | 10X | LR+GLM |
| total length | 951,362,726 | 908,224,480 | 791,708,797 | 877,456,336 | 958,225,486 | 901,238,333 |
| longest 24 sequences assembly fraction | 99.0% | 94.7% | 96.5% | 98.8% | 41.4% | 99.5% |
| scaffold/chromosome N50 | 39,550,354 | 36,400,992 | 33,333,317 | 37,412,490 | 6,260,519 | 41,060,379 |
| contig N50 | 4,101,751 | 3,162,456 | 6,206,245 | 4,268,950 | 12,991 | 6,160,542 |

The genome of *P. schrenkii* was assembled by a hybrid assembly method, which was highly efficient regarding long-read sequencing coverage and read length needed (here only 30-fold / N50 read length: 4.95 kb). *De novo* assembled contigs from short reads were combined with long reads and scaffolded using our CSA-pipeline [38], with the *P. flavescens* as the closest reference genome (Fig. 1: divergence time about 7.1 Mya) for genome comparison and inferring chromosomal-level sequences. Using this approach, we were able to assemble the genome of *P. schrenkii* to similar quality as those obtained for *P. flavescens* and *P. fluviatilis* (94.7% assigned to 24 chr. / contig N50 length: 3.2 Mbp; Table 2). The genome assembly size of *P. schrenkii* was in between the other two *Perca sp.* genomes (877 Mbp < 908 Mbp < 951 Mbp).

**Figure 1:**
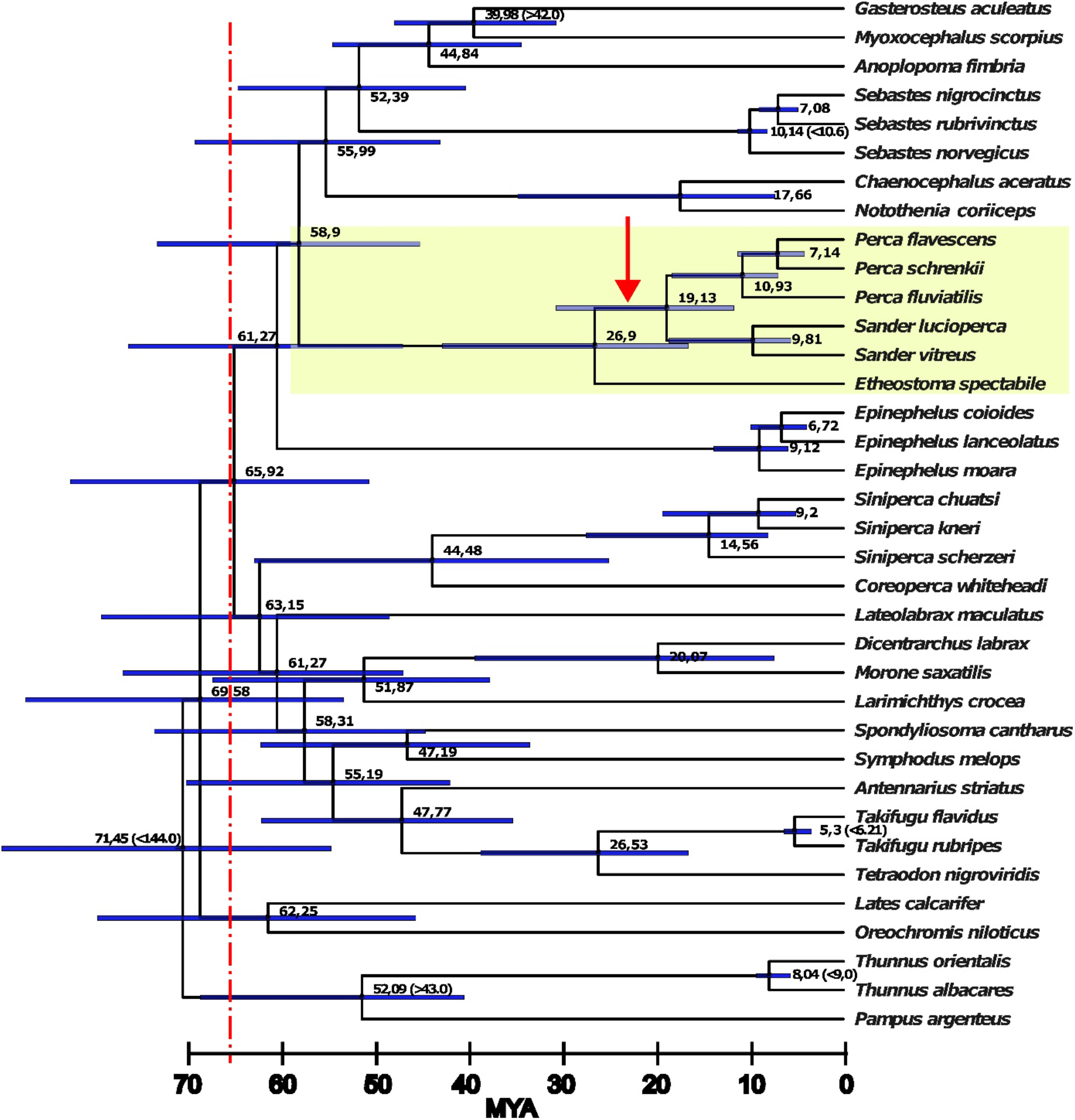
Time calibrated phylogenomic tree constructed from non-coding alignments of 36 *Percomorpha* genome assemblies reveals a massive radiation after the Cretaceous–Paleogene (K–Pg) boundary (66 Mya, dotted red line). The family Percidae is highlighted in yellow. Node numbers depict the median ages in Mya calculated by Mcmctree (values in brackets were taken from www.timetree.org and used for calibration). Blue bars depict the 95% confidence intervals for the node ages. Red arrow indicates duplication event of *amhr2* in *Percidae* (more details in Fig. 2a, *amhr2* gene tree). All branches of the tree obtained 100% support using the SH-aLRT (Shimodaira–Hasegawa like approximate likelihood ratio test) and UFBS (ultra-fast boostrap) tests.

The genome of *Sander vitreus* was assembled from long reads (coverage: 12-fold / N50 read length: 10 kb) and short reads similar to procedures for *P. schrenkii* but we used two reference genomes for chromosomal assembly. First, contigs of *S. lucioperca* (closest relative, divergence time 9.8 Mya) served to order the *S. vitreus* contigs (N50: 6.2 Mb), which improved scaffold N50 significantly (result. N50: 16.8 Mb), then these nearly chromosome-scale scaffolds were ordered according to the *P. flavescens* (div. time 19.1 Mya) Hi-C chromosomes (result. N50: 33.3 Mb / 96.5% assigned to 24 chromosomes). This two-step approach resulted in more consistent results than just using the *S. lucioperca* chromosomal scaffolds, which were generated by genetic linkage mapping. We observed similar genome size differences as in *Perca sp.* between both *Sander* species. The *S. vitreus* genome assembly (791 Mb) is smaller than the one of *S. lucioperca* (901 Mb), thus the North American *Sander* and *Perca* species tend to have smaller genome sizes than their Eurasian relatives (Table 2).

*De novo* repeat analysis showed that 60.1% of the genome assembly size difference between *Sander vitreus* and *S. lucioperca* could be explained by repeat expansion/reduction. Similarly, for *Perca flavescens* and *P. fluviatilis* about 64.8% of the genome size differences were due to repeat expansion/reduction. In both species pairs most repeat expansions/reductions were observed in repeat elements classified as “unknown”. Regarding annotated repeat element classes, L2, DNA, Helitron, CMC-EnSpm, hAT, Rex-Babar, hAT-Charlie and PiggyBac elements expanded the most in both Eurasian species (together adding roughly 20 Mbp to the genomes of *S. lucioperca* and *P. fluviatilis*), while a clear expansion of only a single repeat element, called RTE-BovB, was found in both North American species (adding about 3 and 7 Mbp of sequence to *S. vitreus* and *P. flavescens*, respectively; Suppl. table 1).

The genome of *P. fluviatilis* was annotated by NCBI/GNOMON, which included ample public RNAseq data and protein homology evidence. For *P. schrenkii* and *S. vitreus*, we transferred the NCBI/GNOMON annotations of *P. flavescens* and *S. lucioperca*, respectively. BUSCO analysis (Table 3) revealed values larger than 95.9% for complete BUSCOs (category “C:”) for all annotations. The comparative annotation approach resulted only in small losses (category “M:”) of a few BUSCO genes in the range of 0.4% - 1.1%. In this regard, the *S. vitreus* assembly performed better than the *P. schrenkii* assembly, possibly due to the higher N50 read length of the underlying long-read data.

**Table 3:** BUSCO scoring of annotations of five Percidae genome assemblies. (*P. fluviatilis*, *P. schrenkii*, *S. vitreus* from this study). The comparative mapping of high quality NCBI/GNOMON annotations onto closely related species’ genome assemblies is a cost-effective and fast procedure to annotate new genomes. Abbreviations used: C = complete; S = single copy; D = duplicated; F = fragmented; M = missing.

|  | Species | Annotation type | BUSCO scoring code |
| --- | --- | --- | --- |
| # 1 | *P. fluviatilis* | NCBI/GNOMON | C: 98.5% [S: 95.3%, D: 3.2%], F: 1.0%, M: 0.5%, n: 4584 |
| # 2 | *P. flavescens* | NCBI/GNOMON | C: 99.3% [S: 95.9%, D: 3.4%], F: 0.5%, M: 0.2%, n: 4584 |
| # 3 | *P. schrenkii* | mapping of annot. #2 | C: 95.9% [S: 92.6%, D: 3.3%], F: 2.8%, M: 1.3%, n: 4584 |
| # 4 | *S. lucioperca* | NCBI/GNOMON | C: 99.3% [S: 95.9%, D: 3.4%], F: 0.5%, M: 0.2%, n: 4584 |
| # 5 | *S. vitreus* | mapping of annot. #4 | C: 98.6% [S: 95.5%, D: 3.1%], F: 0.8%, M: 0.6%, n: 4584 |

### Percomorpha phylogenomics and divergence time estimation

To calculate the phylogenetic tree of 36 Percomorpha species and their divergence times, we aligned whole genomes and extracted the non-coding alignments (Fig. 1). The use of non-coding sequences is preferable to calculate difficult-to-resolve phylogenetic trees that occur after massive radiations [60–62]. Our multiple alignment consisted of 6,594,104 nt residues (2,256,299 distinct patterns; 1,652,510 parsimony-informative; 1,136,496 singleton sites; 3,805,098 constant sites) and resulted in a highly-supported tree (raxml-ng and IQtree2 topologies were identical; all IQtree2’s SH-aLRT and ultrafast bootstrapping (UFBS) values were 100). According to the divergence time analysis, most Percomorpha orders emerged after the Cretaceous–Paleogene (K-Pg) boundary about 65.9 Mya ago (CI: 51.3 - 83.6). The lineage leading to the Percidae (represented with species of *Perca*, *Sander*, and *Etheostoma*) emerged about 58.9 Mya (CI: 45.8 - 74.2), and the extant Percidae species analyzed in this study diverged from a last common ancestor (LCA) about 26.9 Mya (CI: 16.8 - 43.4). The *Perca* and *Sander* genera split about 19.1 Mya (CI: 11.8 - 31.1), and *S. vitreus* and *S. lucioperca* splitted at 9.8 Mya (CI: 5.7 - 18.9) similar to the divergence of *P. fluviatilis* from *P. flavescens* and *P. schrenkii* 10.9 Mya (CI: 7.1 - 18.6). The closest *Perca* species are *P. flavescens* and *P. schrenkii* which diverged about 7.1 Mya (CI: 4.2 - 11.4), although today both species are occurring in completely different global ranges.

### The fate of amhr2 genes during evolution of Perca and Sander species

In the genome of *P. flavescens,* two *amhr2* paralogs were previously described, i.e., an autosomal gene, *amhr2a*, present in both males and females on chromosome 04 (Chr04), and a male-specific duplication on the Y-chromosome (Chr09), *amhr2bY* [9]. A similar *amhr2* gene duplication was also found in our male *P. schrenkii* assembly, and sequence homologies and conserved synteny analyses show that these two *P. schrenkii amhr2* genes are orthologs of *P. flavescens amhr2a* and *amhr2bY,* respectively (Fig. 2A and AB). Genotyping of one male and one female also suggests that the *amhr2bY* gene could be male-specific in *P. schrenkii* (Suppl. Fig. 2, Table 4), as described in *P. flavescens* [9]. This potential sex-linkage is also supported by a half coverage of reads in the genomic region containing the *amhr2bY* locus in our male *P. schrenki* genome assembly (Suppl. Fig. 3), in agreement with the hemizygosity of a male-specific region on the Y in species with a XX/XY sex determination system. Alignment of the *P. schrenkii amhr2bY* ortholog shows that its coding sequence (CDS) shares 98% identity with the *P. flavescens amhr2bY* CDS and 95.7 % identity at the protein level the *P. flavescens Amhr2bY*. As in *P. flavescens* [9], the *P. schrenkii amhr2bY* gene encodes a N-terminal-truncated type II receptor protein that lacks most of the cysteine-rich extracellular part of the receptor, which is crucially involved in ligand-binding specificity [63] (Fig. 3).

**Figure 2:**
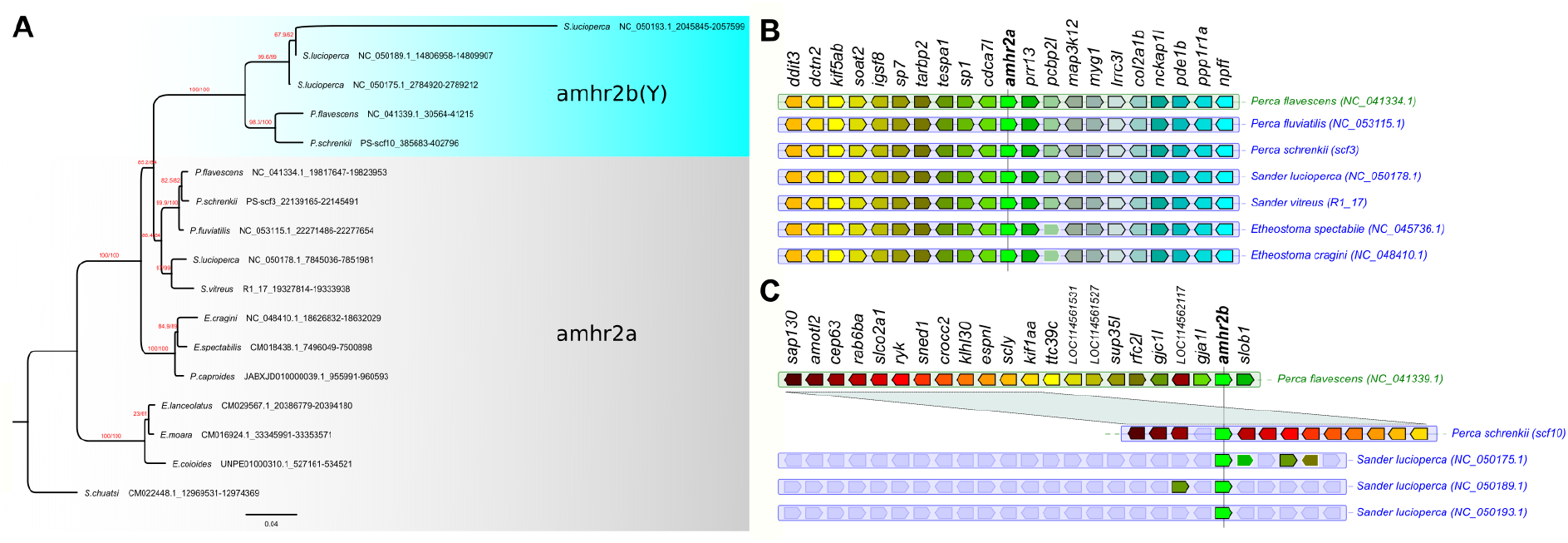
Evolution of *amhr2* genes in Percidae. **A.** Orthologs of *amhr2a* were identified in genome assemblies and their gene tree is consistent with the species tree, which was found by the phylogenomics approach (see Figure 1) The *amhr2b* duplications were only found in the genome assemblies of *P. flavescens* (single copy, male specific), *P. schrenkii* (single copy, potentially male specific) and *S. lucioperca* (three copies, no clear sex linkage) and they clustered together. Thus, *amhr2b* stems from a gene duplication event that occurred at the origin of Percidae (19-27 Mya) and the absence of *amhr2b* in *P. fluviatilis* and *S. vitreus* suggests a secondary loss event in these species. This tree was calculated on codon position 1 and 2 alignments and achieved the best bootstrap support (red numbers: SH-aLRT / UFBS support values) for the split of the *amhr2a* and *b* clades. Trees based on complete CDS, CDS + Introns and amino acid sequences resulted in the same topology albeit with lower bootstrap support for some splits (Suppl. Fig. 8). Numbers after species names depict the coordinates of the respective *amhr2* genes in the genome assemblies. **B & C. Conserved synteny around the *amhr2a* (B) and *amhr2b* (C) loci in some *Percidae* species**. These multiple duplications (in *S. lucioperca*) or the loss of the *amhr2b* genes (in *P. fluviatilis* and *S. vitreus*) emphasize that *amhr2b* may be considered a “jumping” gene locus, which is also supported by conserved synteny analysis.

**Figure 3:**
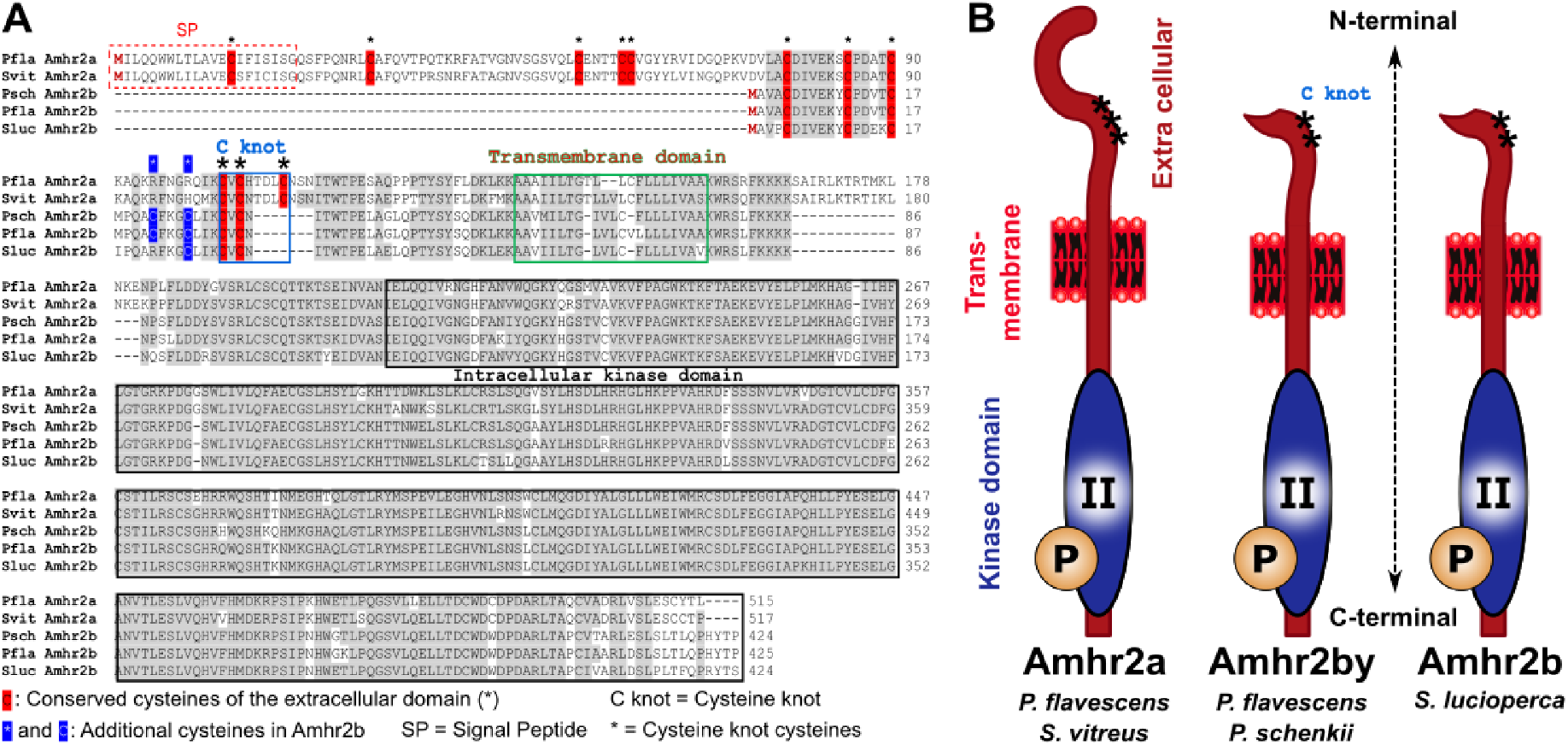
Alignment and structure of Amhr2a and Amhr2b proteins in *Perca flavescens* (Pfla) and *P. schrenkii* (Psch), *Sander lucioperca* (Sluc) and *S. vitreus* (Svit). **A**. Alignment of some Percidae Amhr2a (*P fluviatilis* and *S. vitreus*) and Amhr2b (*P. schrenkii*, *P. flavescens* and *S. lucioperca*) proteins. **B**. Schematic structure of Amhr2a and Amhr2b proteins.

**Table 4:** Sex-linkage of different sex-markers in *Perca fluviatilis* and *Perca schrenkii*. Associations between each sex-specific marker and sex phenotypes are provided for both males and females (number of positive individuals / total number of individuals) along with the p-value of association with sex that was calculated for each species and population based on the Pearson’s Chi-square test with Yates’ continuity correction. ns = not statistically significant.

| Species | Population | marker | males | females | p value |
| --- | --- | --- | --- | --- | --- |
| *Perca schrenkii* | Alakol Lake, Kazakhstan | *amhr2bY* | 1/1 | 0/1 | ns |
| *Perca fluviatilis* | Mueggelsee Lake Germany | *hsdl1* | 10/10 | 0/9 | 9.667e-05 |
| *Perca fluviatilis* | Lucas Perche, France | SNV1 | 48/48 | 0/47 | < 2.2e-16 |
| *Perca fluviatilis* | Kortowskie Lake, Poland | SNV1 | 17/17 | 0/20 | 8.83e-09 |

In contrast, in our *P. fluviatilis* male genome assembly, only one copy of *amhr2* could be identified and sequence homologies and conserved synteny analysis (Fig. 2A and 2B) shows that this *amhr2* gene is the ortholog of *P. flavescens* and *P. schrenkii amhr2a*. In addition, PCR with primers designed to amplify both *amhr2a* and *amhr2bY* from *P. flavescens* and *P. schrenkii* did not show any sex-differences in *P. fluviatilis*. Altogether, these results support the absence of an a*mhr2b* gene in *P. fluviatilis*.

In the *Sander lucioperca* male genome assembly, four copies of *Amhr2* were detected. Using the same primers as those used for the amplification of both *amhr2a* and *amhr2bY* in *P. flavescens* and *P. schrenkii*, PCR genotyping on males and females of *S. lucioperca* produced complex amplification patterns with multiple bands and no visible association with sex. In the publicly available genome assembly of *Sander vitreus,* sequence homologies and /or conserved synteny analysis (Fig. 2A and 2B) allowed the identification of a single autosomal *amhr2a* gene.

A phylogenetic analysis of the sequences with similarity to *amhr2* in *Perca* and *Sander* (Fig. 2A and 2B) shows that the different *amhr2* genes likely originated from a gene duplication event that happened in the branch leading to the last common ancestor of these species, dated around 19-27 Mya. Since that time, the *amhr2bY* gene has been conserved in *P. schrenkii* and *P. flavescens*, lost in *P. fluviatilis and S. vitreus,* and amplified in *S. lucioperca*.

### Evolution of sex determination in P. fluviatilis

Because *P. fluviatilis* sex determination does not rely on an *amhr2* duplication like what has been found in *P. flavescens* [9] and *P. schrenkii* (this study), we used genome-wide approaches to better characterize its sex-determination system. RAD-Seq analysis on 35 males and 35 females of *P. fluviatilis*, carried out with a minimum read depth of one, allowed the characterization of a single significant sex marker (Suppl. Fig. 4). This 94 bp RAD sequence matches (Blast Identities: 93/95%) a portion of *P. fluviatilis* chromosome 18 (GENO_Pfluv_1.0, Chr18: 27656212 - 27656305). This RAD-Seq analysis suggests that Chr18 could be the *P. fluviatilis* sex chromosome, and that its sex-determining region could be very small because we only detected a single significant sex-linked RAD-sequence. To get a better characterization of the *P. fluviatilis* sex chromosome and sex-determining region, we then used Pool-Sequencing (Pool-Seq) to re-analyze the same samples used for RAD-Seq by pooling together DNA from the males in one pool and DNA from females in a second pool. Using these Pool-Seq datasets, we identified a small 100 kb region on *P. fluviatilis* Chr18 with a high density of male-specific SNVs (Fig. 4), confirming the RAD-Seq hypothesis that Chr18 is the sex chromosome in that species. No male-specific duplication / insertion event was found in this sex-determining region on Chr18, which contains six annotated genes (Fig. 4D). These genes encode a protein of unknown function (*C18h1orf198*), three gap-junction proteins (*Cx32.2, Gja13.2* and *Cx32.7*), a protein annotated as inactive hydroxysteroid dehydrogenase-like protein (*Hsdl1*), and a protein known as protein broad-minded or Tcb1 domain family member 32 protein (*Tbc1d32*). Three of these six annotated genes, i.e., *c18h1orf198*, *hsdl1* and *tbc1d32* display a higher expression in testis than in the ovary (Suppl. Fig. 5).

**Figure 4:**
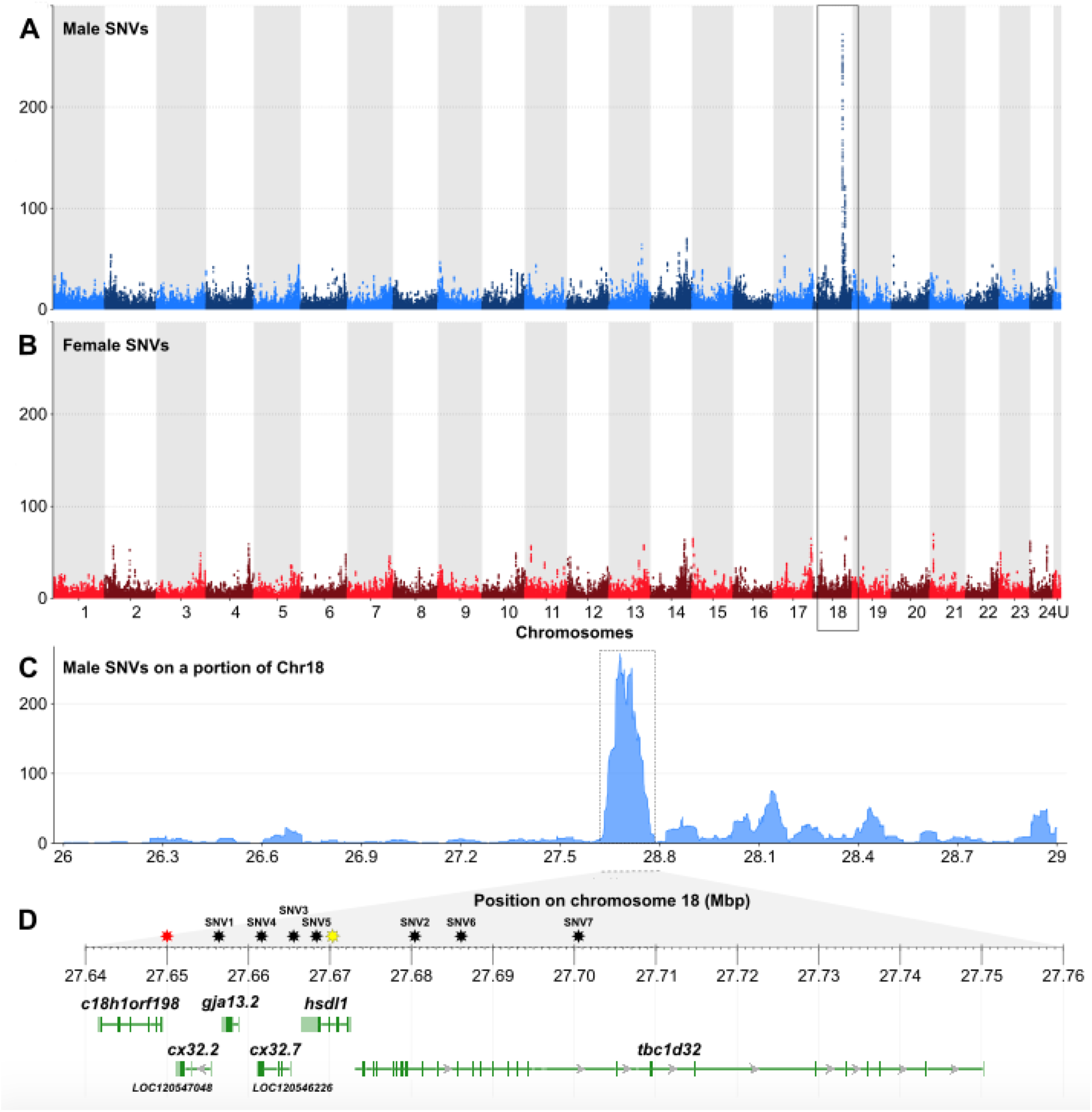
Chromosome 18 is the sex chromosome of *Perca fluviatilis*. (**A**, **B**) Genome-wide Manhattan-plots of (**A**) male- and (**B**) female- specific single-nucleotide variations (SNVs), showing that chromosome 18 (Chr18) contains a 100 kb region, enriched for male-specific SNVs. Male- and female- specific SNVs are represented as dots (total of SNVs per 50 kb window size) of alternating colors on adjacent chromosomes. (**C**) Zoomed view of the male-specific SNVs on the sex-biased region of Chr18 with its gene annotation content (**D**): *c18h1orf198* = c18h1orf198 homolog ; *cx32.2* (LOC120547048) = gap junction Cx32.2 protein-like ; *gja13.2* = gap junction protein alpha 13.2 ; *cx32.7* (LOC120546226) = gap junction Cx32.7 protein-like ; *hsdl1* = inactive hydroxysteroid dehydrogenase-like protein 1 ; *tbc1d32* = tcb1 domain family member 32 (also known as protein broad-minded). Stars over the ruler of panel D are the locations of the single male-specific RAD maker (red star) and of the SNVs used for designing KASPar assays (black stars; SNV1-7). The location of the sex-specific intronic indel inside the *hsdl1*-gene used for sex specific PCR is shown by a yellow star

To provide a better support for the sex-linkage of the male-specific variants found within this sex determining region on Chr18, we designed different types of genotyping assays (classical PCR and KASpar) that have been applied to different *P. fluviatilis* individuals which were phenotypically sexed with confidence. A classical PCR-assay was first developed based on the detection of a Y-allele-specific 27 bp deletion in the third intron of the *P. fluviatilis hsdl1* gene, (Suppl. Fig. 6, Table 4) and this assay successfully identified all males of a Lake Mueggelsee from Germany (10 males and 9 females; p-value of association with sex = 9.667e-05). In addition, KASpar allele-specific PCR-assays were developed based on seven single nucleotide sex-specific variants, located at different positions within the Chr18 sex-determining region of P. *fluviatilis* (Fig. 3D). Tests of 48 males and 48 females showed that of seven KASpar allele-specific PCR-assays, five resulted in a high proportion of correctly-genotyped individuals with males being heterozygote and females being homozygote (>95%). Two of the targeted SNVs (SNV1 and SNV3) that displayed 100% sex-linkage accuracy (Suppl. Fig. 7, Tables 4 and 5). Sex-linkage of SNV1 was then checked on a wild-type population from Kortowskie Lake in Poland for which the association of male phenotype and SNV1 heterozygosity was also complete (17 males and 20 females; p-value = 8.83e-09, Table 4).

**Table 5:** KASpar allele-specific PCR assays on seven single sex-specific nucleotide variations (SNV ID#) in P. *fluviatilis*. Numbers of homozygote (Ho), heterozygote (He) and uncalled genotypes (U). N = Total numbers of genotyped individuals (M/F: Males/females), %N = percentage of genotyped individuals, % As = percentage of correctly assigned genotypes i.e., male heterozygotes and female homozygotes. The p-value of association with sex was calculated for each SNV based on the Pearson’s Chi-square test with Yates’ continuity correction scoring heterozygote males and homozygote females as positives.

| SNV ID# | N (M/F) | % N | M Ho/He/U | F Ho/He/U | % As | p value |
| --- | --- | --- | --- | --- | --- | --- |
| SNV1 | 95 (48/47) | 99,0 | 0/48/0 | 47/0/0 | 100,0 | < 2.2e-16 |
| SNV2 | 96 (48/48) | 100,0 | 8/37/3 | 34/2/12 | 74,0 | 3.175e-11 |
| SNV3 | 93 (45/48) | 96,9 | 0/45/0 | 48/0/0 | 100,0 | < 2.2e-16 |
| SNV4 | 96 (48/48) | 100,0 | 0/46/2 | 46/0/2 | 95,8 | < 2.2e-16 |
| SNV5 | 92 (44/48) | 95,8 | 0/42/2 | 46/0/2 | 95,7 | < 2.2e-16 |
| SNV6 | 93 (48/45) | 96,9 | 1/47/0 | 25/2/18 | 77,4 | 1.964e-14 |
| SNV7 | 95 (47/48) | 99,0 | 1/46/0 | 45/3/0 | 95,8 | < 2.2e-16 |

## DISCUSSION

The Percidae family is represented by 239 species and 11 genera. The genera *Perca* and *Sander* are especially important for aquaculture and fisheries. By providing new genome sequence assemblies for *Perca fluviatilis*, *P. schrenckii* and *Sander vitreus*, we provide for the first-time access to all economically important species of the *Percidae* at the DNA-level. Assembly statistics for these three new genome sequence assemblies are reference grade with N50 continuity in the megabase range and chromosomal length scaffolds, obtained either by Hi-C scaffolding or by conserved synteny analysis involving the closest relatives. Importantly completeness on the gene-level is significantly higher than in a short-read based draft genome for *P. fluviatilis*, published earlier [64], allowing much stronger conclusions to be drawn regarding the presence or absence of possible sex-determining genes.

### The origin of Percidae

A phylogenomic approach using aligned non-coding sequences of 36 genomes resulted in a highly-supported tree showing a rapid radiation of fish families. It has recently been shown that phylogenomics based on non-coding sequences may be more reliable at resolving difficult-to-resolve radiations in species trees (i.e. in Aves). According to our time calibration, many taxonomic orders related to the Percidae emerged shortly after the Cretaceous-Paleogene (K-Pg) mass extinction event, about 66 Mya, and gave rise to Percidae about 59 Mya. This is in contrast to many older studies, which have, for example, dated the split of *Perca ssp.* and *Gasterosteus ssp*. back to the Cretaceous (73-165 Mya; 18 of 23 studies listed at www.timetree.org). Similar patterns in rapid radiations have been observed in the avian tree of life and have likewise been attributed to the K-Pg mass extinction [61,62]. In context, it has been argued that the so-called “Lilliput effect”, which describes the selection in favor of species with small body sizes and fast generation times after mass extinction events, can lead to an increase in substitution rates and results in overestimations of node-ages for molecular clocks [65].

### The evolution of sex determination in Perca and Sander species

#### Evolution and turnover of Amhr2

Sex determination systems with MSD evolved from duplications of the *amh* [10–17] or *amhr2* [9,18–21] genes have now been characterized in many fish species, all with a male-heterogametic system (XX/XY). In addition, the fact that *Amh* in monotremes [66], or *Amhr2* in some lizards [67] are Y-linked also makes them strong MSD gene candidates in other vertebrate species. In Percidae, sex determination has only been explored in some species of *Perca* [8,9], and *Amhr2* has been characterized as a potential MSD gene in yellow perch, *P. flavescens* [9]. Our results suggest that this duplication of *amhr2* in *P. flavescens* is also shared by *P. schrenkii* and *S. lucioperca*, implying an origin of duplication in their last common ancestor, dated around 19-27 Mya. However, the fate of this duplication seems to be complex - with multiple duplications/insertions on different chromosomes with no clear sex-linkage in *S. lucioperca*, a secondary loss in *P. fluviatilis* and *S. vitreus*, contrasting with a single potentially sex-linked duplication/insertion in *P. schrenkii* and in *P. flavescens*. This finding suggests that the shared ancestral *amhr2b*-duplicated locus (Fig. 2A) might be a jumping locus that has been moving around during its evolution as found for the *sdY* MSD jumping sex locus in salmonids [68,69]. Additional evidence that these *amhr2b* genes originated a single ancestor also rely on the fact that the *Amhr2b* proteins of *P. flavescens* [9], *P. schrenkii* and *S. lucioperca* share a similar gene structure with an N-terminal truncation that results in the absence of the cysteine-rich extracellular part of the receptor. This part of the receptor is known to be crucial for ligand binding [70]. A similar N-terminal truncation of a duplicated *amhr2* was also described in catfishes from the Pangasidae family, where this truncation had been hypothetically linked to a potentially new sex-determination function that lacks ligand dependency [18]. In the genus *Sander*, the situation might be similar to that in *Perca* regarding the changes of the sex-determination systems between species. In *S. lucioperca*, *amhr2b* might still serve as the MSD-gene, but the several recent *amhr2b* duplications have complicated our analysis so far. Similar to *P. fluviatilis*, *S. vitreus* has lost *amhr2b* and likely another factor took over as a potential MSD-gene.

#### A new sex-specific locus in P. fluviatilis

The fact that *P. fluviatilis* sex does not rely on an *amhr2*-duplication like *P. flavescens* [9] and *P. schrenkii* do, indicates that *P. fluviatilis* evolved a completely different and new MSD-gene. Our results also show that this sex locus on *P. fluviatilis* Chr18 is very small compared to what is observed in many fish species, with a potentially non-recombining size around 100 kb. This locus however is not the smallest SD-locus described in teleost fish: in the pufferfish, *Takifugu rubripes*, the sex locus is limited to a few SNPs that differentiate *amhr2* alleles on the X- and Y -chromosomes [19]. Because we did not find any sign of a sex chromosome-specific duplication/insertion event in the *P. fluviatilis* SDR, this sex locus seems to result from pure allelic diversification and is thus in contrast to *P. flavescens* [9] and probably also *P. schrenkii* (this study). The *P. fluviatilis* sex specific-region on Chr18 contains six annotated genes, which encode a protein of unknown function (*C18h1orf198*), three gap-junction proteins (*Cx32.2, Gja13.2* and *Cx32.7*), a protein annotated as inactive hydroxysteroid dehydrogenase-like protein (*Hsdl1*) and a protein known as protein broad-minded or Tcb1 domain family member 32 protein (*Tbc1d32*). Of these genes, *hsdl1* and *tcb1d32* are interesting potential MSD candidates in *P. fluviatilis,* based on their potential functions and the fact that both display a higher testicular expression compared to the ovary. The Hsdl1 protein is indeed annotated as “inactive” [71], but this annotation only refers to its lack of enzymatic activity against substrates so far tested, leaving other potential functional roles for a protein that is highly conserved in vertebrates [71]. In *Epinephelus coioides*, *hsdl1* has been shown to be differentially expressed during female-to-male sex-reversal, and its expression profile clustered with *hsd17b1* [72], which plays a central role in converting sex steroids and has recently been identified as a potential MSD gene in different species with a female heterogametic (ZZ/ZW) sex determination system, like in oyster pompano, *Trachinotus anak* [73] and different amberjack species [74,75]. The *Tbc1d32* protein has been shown to be required for a high Sonic hedgehog (*Shh*) signaling in the mouse neural tube [76]. Given the role of *Shh* signaling downstream of steroidogenic factor 1 (*nr5a1*) for the proper steroidogenic lineage fate [77] and the importance of steroids in gonadal sex differentiation [78,79], *tbc1d32* would be also an interesting candidate as a potential MSD gene in *Perca fluviatilis*.

## CONCLUSIONS

Our study shows that *Percidae* exhibit a remarkable high variation in sex-determination mechanisms. This variation is connected to deletion or amplification of *amhr2bY*, which if lost in certain species (like *Perca fluviatilis* or *Sander vitreus*) should cause re-wiring of the sex determining pathways and result in the rise of new SD-systems. The mechanisms behind a “jumping” *amhr2bY* expansion or loss and which genes replace it as the MSD remain to be elucidated. The new *Percidae* reference sequence assemblies presented here and the highly reliable sex markers developed for *P. fluviatilis* can now be applied for sex genotyping in basic science as well in aquaculture.

## DATA AVAILABILITY

All genome assemblies and raw sequence datasets have been submitted to NCBI/GENBANK under the bioproject accessions: PRJNA549142, PRJNA637487, PRJNA808842 (*P. fluviatilis*, *P. schrenkii*, *S. vitreus*, respectively).

## BENEFIT-SHARING STATEMENT

A research collaboration was developed with scientists from the countries providing genetic samples (PE and WL in USA, DZ in Poland and ST in Russia), all collaborators are included as co-authors, the results of research have been shared with the provider communities, and the research addresses a priority concern, in this case the conservation of organisms being studied. More broadly, our group is committed to international scientific partnerships, as well as institutional capacity building.

## ACKNOWLEDGEMENTS

We kindly acknowledge the NCBI/Genbank team for providing a GNOMON annotation of the *P. fluviatilis* genome. This work was funded by the German Research Foundation (DFG) “Eigene Stelle” grant KU 3596/1-1; project number: 324050651, by the Agence Nationale pour la Recherche (ANR) / DFG PhyloSex project (ANR-13-ISV7-0005), the CRB-Anim “Centre de Ressources Biologiques pour les animaux domestiques” project PERCH’SEX, the FEAMP “Fonds européen pour les affaires maritimes et la pêche” project SEX’NPERCH, NIH (R35 GM139635), and NSF (2232891). The GeT-PlaGe and MGX sequencing facilities were supported by France Génomique National infrastructure as part of the “Investissement d’avenir” program managed by (ANR-10-INBS-09). We thank Dr. Tatjana Dujsebayeva, Salamat Karlybai and his colleague for help during field work in Kazakhstan. We thank Eva Kreuz and Wibke Kleiner for technical assistance. We thank Matt Faust (Ohio, DNR) for help with samples of *S. vitreus*.

## AUTHOR CONTRIBUTIONS

YG and HK designed the project. PE, WL provided sexed samples of *S. vitreus*. MW, BS, TL, ST, SV, DZ collected and sexed the *P. fluviatilis* samples. MS in collaboration with ST did field work in Kazakhstan, collected and sexed the tissue samples of *P. schrenkii*. EJ, MW, CR, CI, HP and HK extracted the gDNA, made the genomic libraries and sequenced them. CeC, ChK, MZ, MW, ClK, RF, AH, HK and YG processed the genome assemblies and/or analyzed the results. CP, LJ, CaC processed and analyzed the sex genotyping tests. HK and YG wrote the manuscript with inputs from all other coauthors. JHP, CD, HK and YG, supervised the project administration and raised funding. All the authors read and approved the final manuscript.

## COMPETING INTERESTS

All authors declare no competing interests.

## Supplementary figures

**Supplementary Figure 1:**
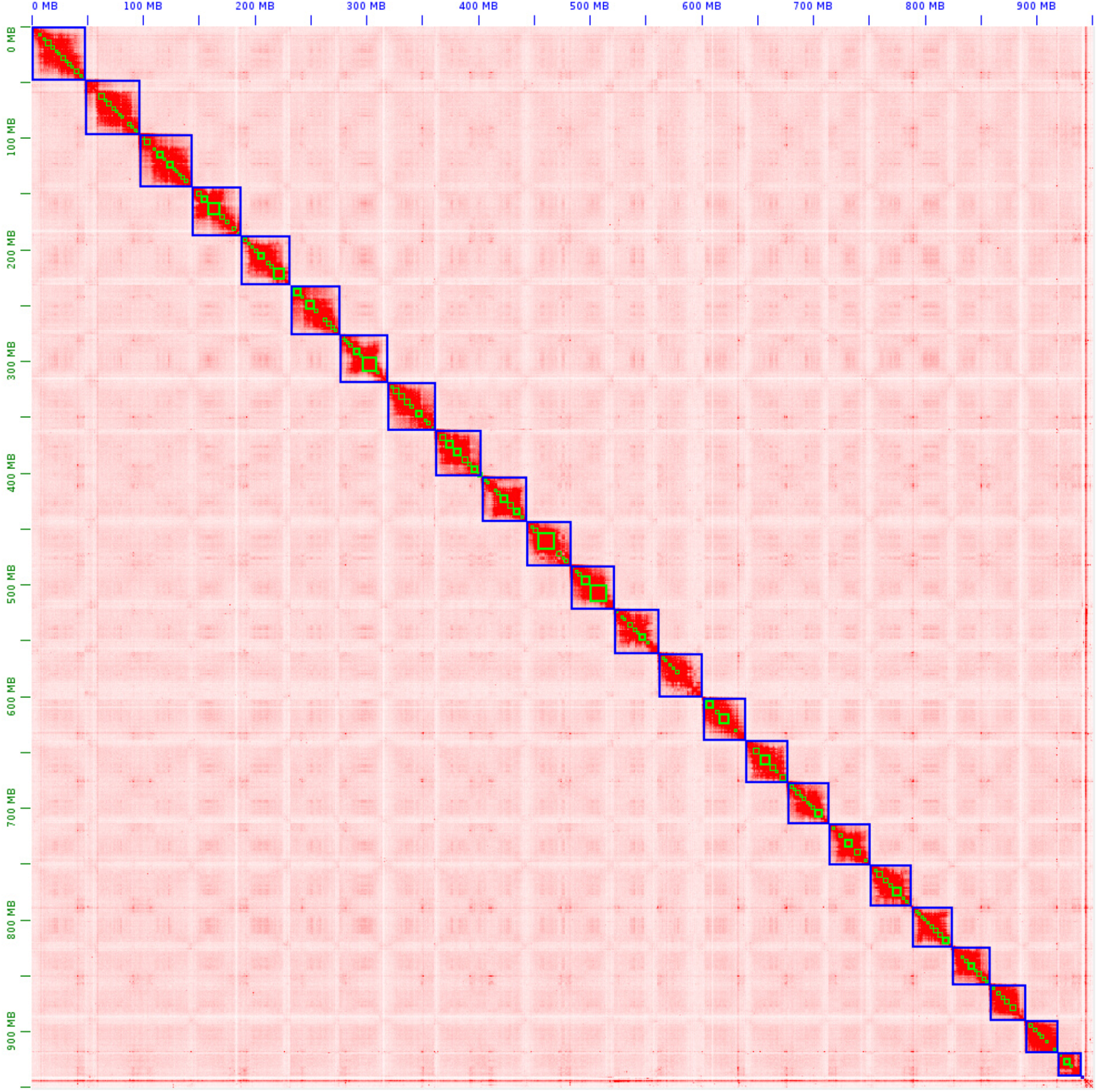
Hi-C map of *P. fluviatilis* genome.

**Supplementary Figure 2:**
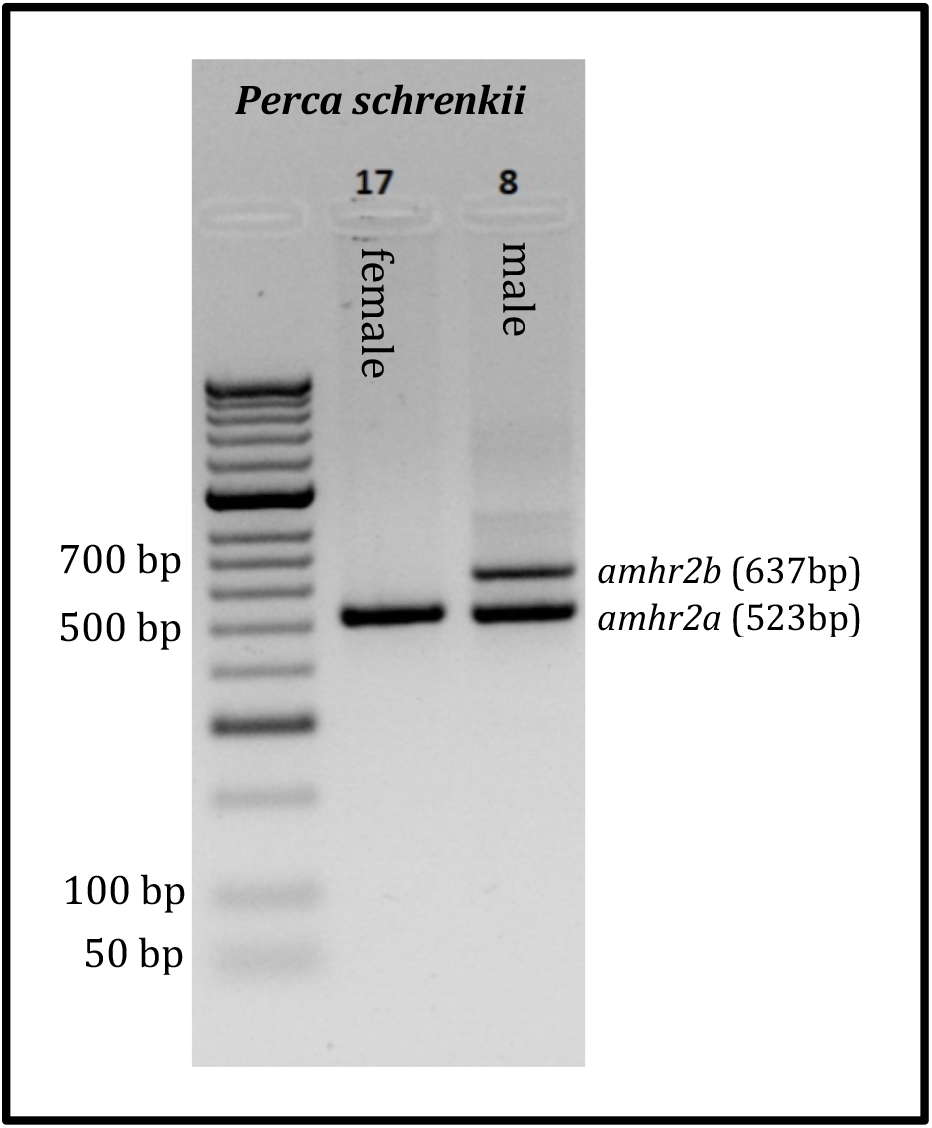
A sex-specific 637 bp PCR product amplifies in *P. schrenkii* male (sample id = 8), while it is absent in female (sample id = 17). The corresponding primer pair also works for sexing of *P. flavescens*.

**Supplementary Figure 3:**
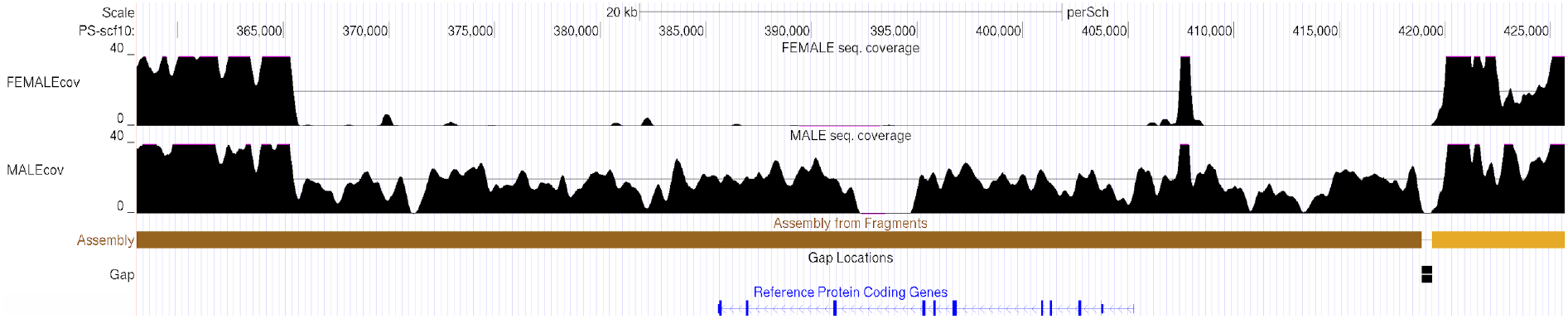
Sex-specific sequencing coverage of *amhr2b* locus in *Perca schrenkii* in region PS-scf10 / CM046795.1: 366,025-418,970. A female and a male genome were sequenced to approximately 40x coverage using short-read sequencing. After filtering for unique mapping reads (mapping quality 60), a clear coverage difference between females and males is visible. The ∼53 kbp region has virtually no coverage in females. In contrast males exhibit haploid coverage (about 20x), which is in accordance with a X/Y SD system.

**Supplementary Figure 4:**
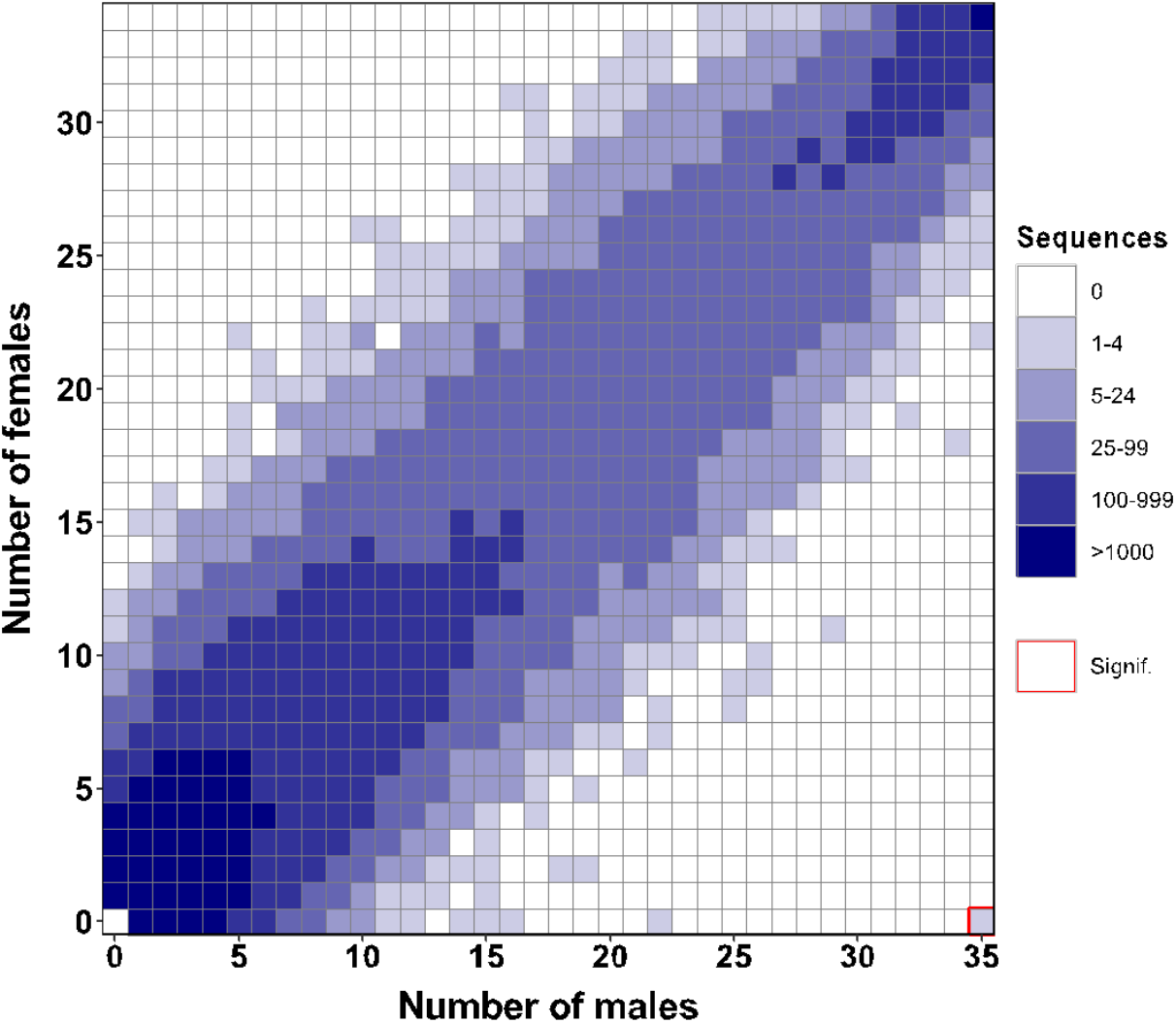
A single RAD sex-specific marker is significantly associated with male sex in *P. fluviatilis*. Tile plot of the distribution of RADSex markers between *Perca fluviatilis* males (horizontal axis) and females (vertical axis) with a minimum read depth of 1 (d = 1). Color intensity (see color legend on the right) indicates the number of markers present for each of the corresponding number of males and females. A single significant marker at the lower right of the grid was present in all 35 males and absent from all 34 females and is boxed with a red border (Chi-squared test, p < .05 after Bonferroni correction).

**Supplementary Figure 5:**
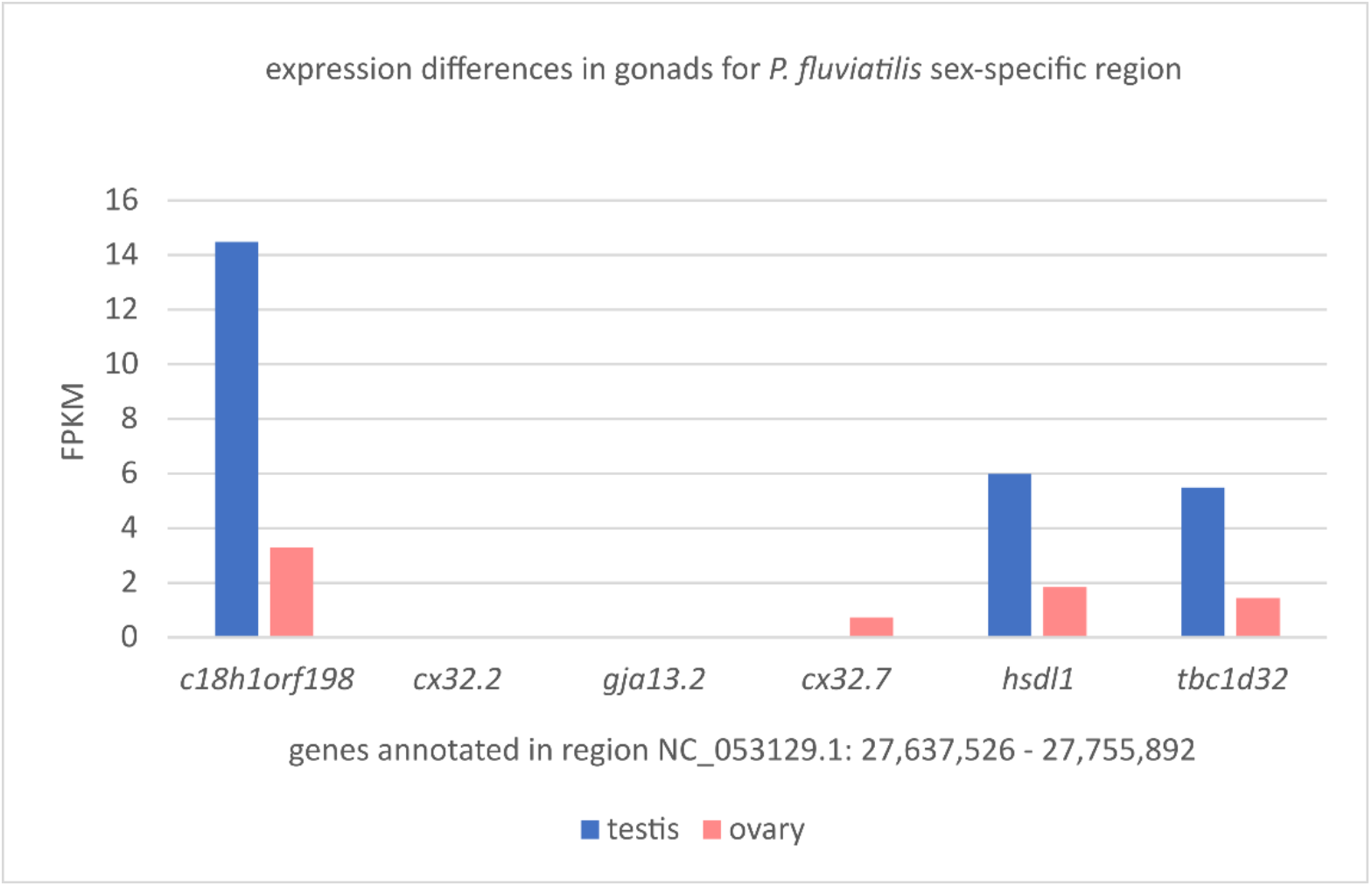
Expression of *hsdl1* and neighboring genes in public RNAseq datasets (testis: SRR14461526, ovary: SRR14461527; age of both sampled individuals 9 month). Here *hsdl1* expression in testis is 3.25-fold higher than in ovary, for *tbc1d32* and *c18h1orf198* the ratio is 3.83 and 4.41, respectively.

**Supplementary Figure 6:**
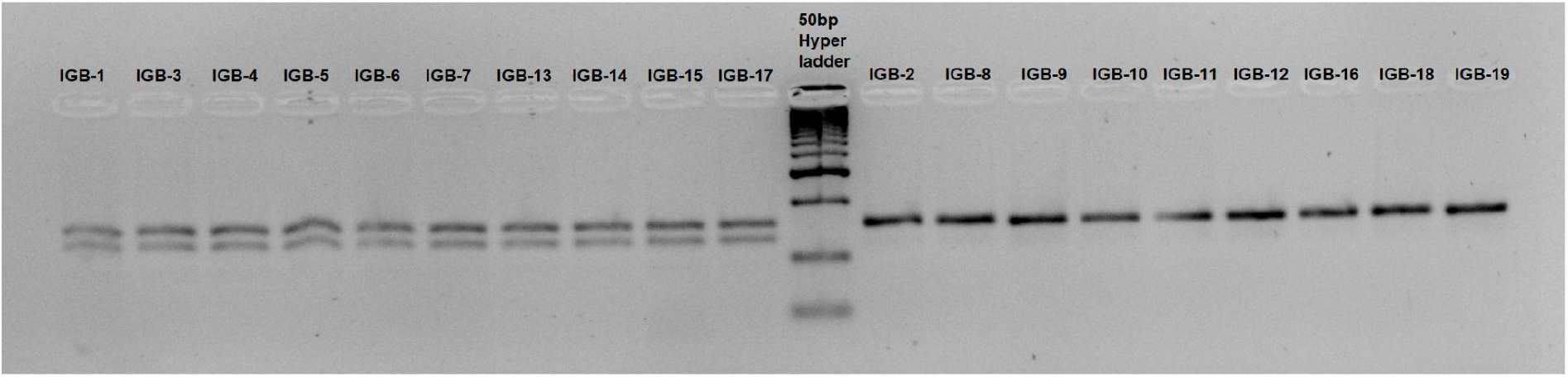
Sexing of *P. fluviatilis* using a 27 bp male specific deletion in Intron 3 of the *hsdl1* gene (10 males (left) and 9 females (right) from Lake Mueggelsee, Berlin. The simultaneous amplification in both males and females of the X allele without the 27 bp deletion provides an internal control for this PCR. All XY male samples (N = 10) produce two amplicons due to the small size difference of the X and Y amplified alleles, and all XX females (N = 9) produced only the larger X amplicon. This *hsdl1* intronic indel variant is located near the variant SNV5 (distance < 1.5 kbp).

**Supplementary Figure 7:**
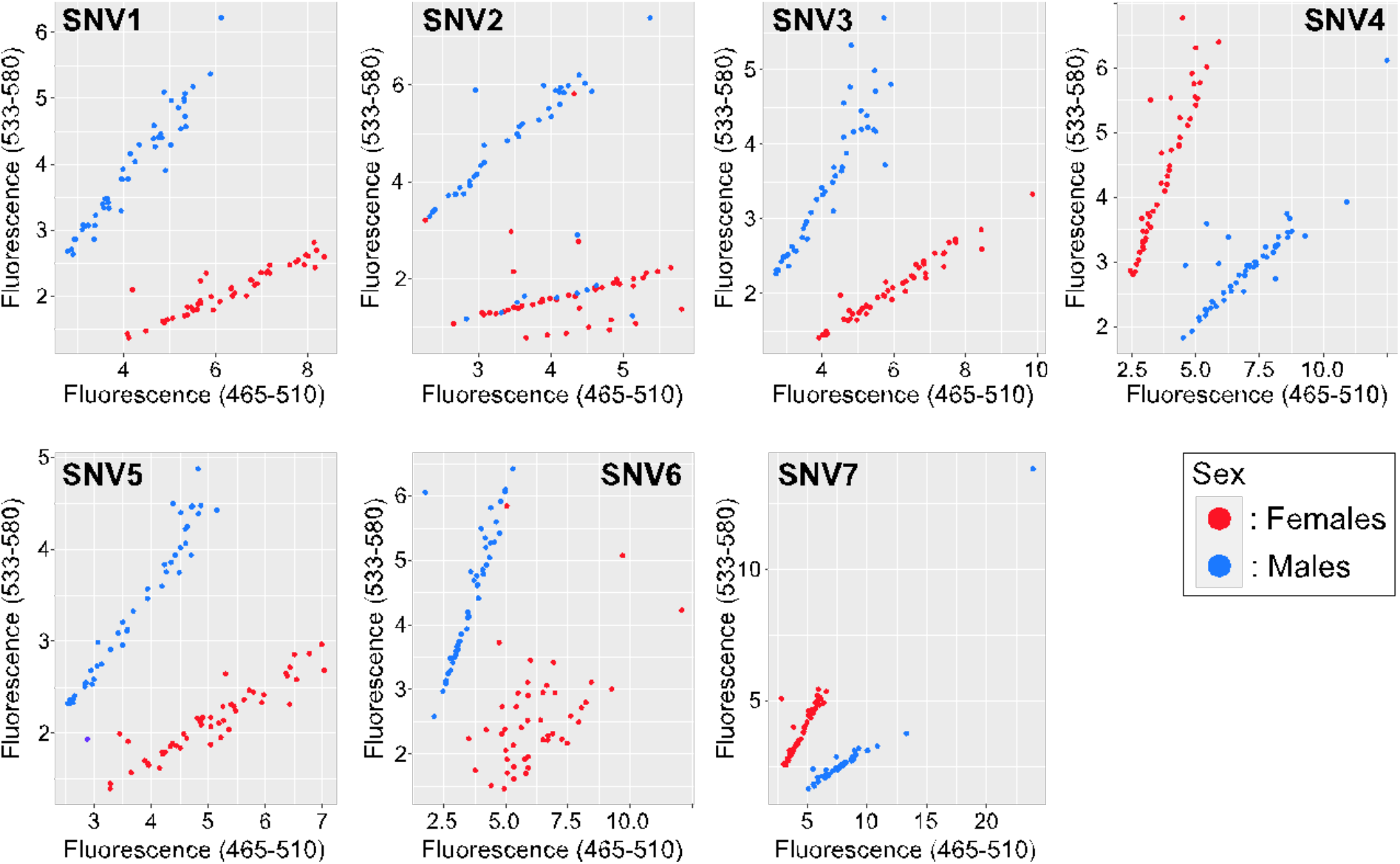
KASpar allele-specific PCR assays on seven single sex-specific nucleotide variations (SNV ID#) in P. *fluviatilis*. For each Single Nucleotide Variation (SNV), primer AL1 was coupled to FAM fluorescent dye and primer AL2 was coupled to VIC fluorescent dye and the end-point fluorescence of these two fluorescent dyes was respectively on the x- and y- axes. Male individuals are represented by blue dots and females by red dots. Primers used for analysis can be found in Table 1.

**Supplementary Figure 8:**
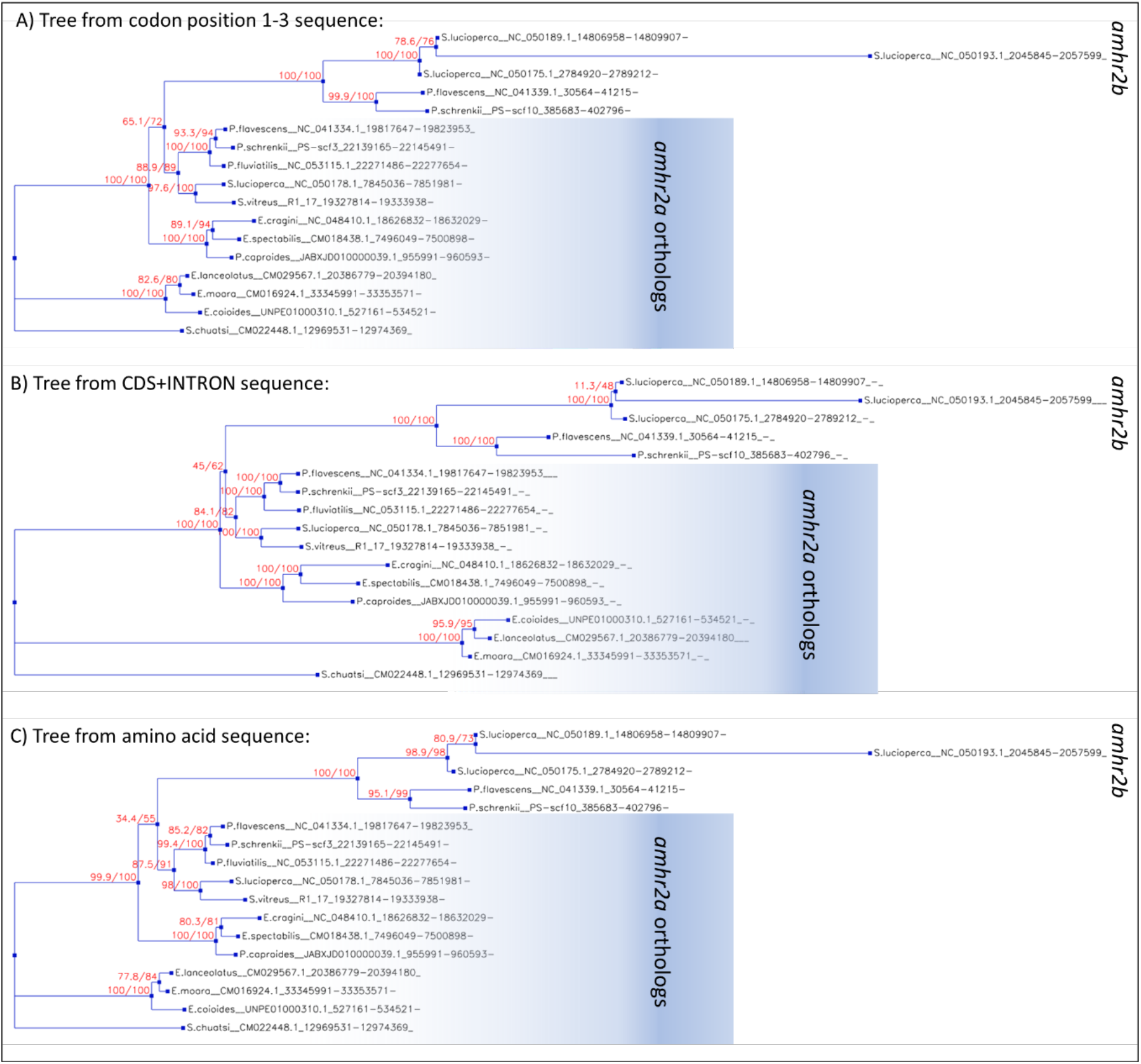
Additional gene trees for *amhr2*. A) Tree calculated from coding sequence. B) Tree calculated from coding plus intron sequence. C) Tree calculated from amino acid sequence. All trees share the same topology but differ in support values for some splits (SH-aLRT and UFBS tests).

**Supplementary Table 1:** Annotated repeats in *Perca sp.* and *Sander sp.* genomes (RepeatModeler *de novo* analysis). Repeat elements mentioned in the manuscript have grey highlighting. The class “DNA” is assigned to repeat elements that harbor signals of transposases, but miss further signals to classify them with more detail.

|  | *P. fluviatilis* | *P. flavescens* | *P. schrenkii* | *S. lucioperca* | *S. vitreus* |
| --- | --- | --- | --- | --- | --- |
| **length (no gaps)** | 951.053.269 | 877.041.836 | 893.440.234 | 901.028.039 | 785.350.625 |
| **Repeat class** | **bp masked** | **bp masked** | **bp masked** | **bp masked** | **bp masked** |
| DNA | **22.711.613** | 20.520.179 | **23.132.743** | **19.957.261** | 15.428.015 |
| Academ | 2.340.071 | 1.619.886 | 1.842.836 | 2.412.073 | 1.790.843 |
| CMC-Chapaev-3 | 230.562 | 0 | 0 | 0 | 0 |
| CMC-EnSpm | **9.118.071** | 7.588.794 | **4.039.674** | **5.928.087** | 3.520.268 |
| Crypton | 841.031 | 547.307 | 748.707 | 772.498 | 402.911 |
| Crypton-V | 11.212 | 0 | 0 | 0 | 0 |
| Dada | 0 | 33.030 | 206.119 | 0 | 0 |
| Ginger | 0 | 15.288 | 0 | 0 | 0 |
| IS3EU | 1.282.033 | 531.773 | 496.067 | 288.665 | 509.614 |
| Kolobok-Hydra | 0 | 648.047 | 0 | 0 | 70.720 |
| Kolobok-T2 | 1.252.506 | 1.976.511 | 2.668.034 | 1.586.132 | 1.222.046 |
| MULE-MuDR | 251.959 | 1.716.319 | 441.114 | 266.625 | 27.741 |
| Maverick | 2.187.699 | 1.402.756 | 266.487 | 1.342.717 | 0 |
| Merlin | 0 | 0 | 0 | 122.531 | 94.494 |
| Novosib | 0 | 0 | 0 | 0 | 82.199 |
| P | 467.516 | 442.149 | 411.688 | 765.209 | 555.234 |
| PIF-Harbinger | 12.989.790 | 12.205.497 | 12.181.221 | 12.830.055 | 8.037.073 |
| PIF-ISL2EU | 0 | 235.589 | 464.262 | 1.597.340 | 914.106 |
| PiggyBac | **3.844.887** | 2.011.288 | **2.742.549** | **3.776.961** | 2.299.194 |
| Sola | 1.885.532 | 229.560 | 163.435 | 77.871 | 0 |
| TcMar | 17.070 | 135.873 | 0 | 185.840 | 137.413 |
| TcMar-Fot1 | 0 | 30.498 | 0 | 44.754 | 37.051 |
| TcMar-ISRm11 | 4.570.634 | 3.171.866 | 2.722.899 | 2.158.133 | 1.830.923 |
| TcMar-Stowaway | 220.092 | 98.992 | 114.404 | 65.697 | 0 |
| TcMar-Tc1 | 15.259.371 | 12.137.252 | 12.302.578 | 6.557.230 | 10.225.920 |
| TcMar-Tc2 | 627.258 | 465.068 | 299.455 | 227.802 | 468.716 |
| Zator | 0 | 119.597 | 0 | 0 | 0 |
| Zisupton | 30.281 | 0 | 127.852 | 0 | 104.202 |
| Zisupton-hAT-hybrid | 0 | 0 | 0 | 0 | 339.346 |
| hAT | **9.251.059** | 7.244.672 | **9.113.922** | **8.460.295** | 6.356.798 |
| hAT-Ac | 26.161.464 | 27.815.352 | 29.153.525 | 30.578.355 | 27.361.032 |
| hAT-Blackjack | 1.252.051 | 459.611 | 338.327 | 514.383 | 437.930 |
| hAT-Charlie | **7.578.170** | 5.645.728 | **8.825.006** | **5.557.301** | 3.893.733 |
| hAT-Tip100 | 3.892.297 | 2.083.169 | 5.060.148 | 2.183.415 | 2.136.454 |
| hAT-Tol2 | 7.383.933 | 7.130.470 | 10.507.431 | 6.788.875 | 4.948.115 |
| hAT-hAT5 | 1.008.628 | 1.141.196 | 1.066.203 | 873.439 | 560.575 |
| hAT-hAT6 | 153.938 | 129.767 | 0 | 0 | 37.338 |
| hAT-hobo | 693.242 | 589.992 | 435.263 | 245.056 | 840.680 |
| LINE | 253.345 | 0 | 0 | 135.963 | 162.701 |
| CR1 | 61.557 | 52.996 | 46.593 | 68.770 | 58.461 |
| Dong-R4 | 0 | 161.938 | 0 | 0 | 0 |
| I | 902.480 | 1.009.950 | 1.721.862 | 451.473 | 448.921 |
| I-Nimb | 187.474 | 66.650 | 179.364 | 0 | 89.737 |
| Jockey | 1.281.097 | 0 | 0 | 249.376 | 0 |
| L1 | 3.161.932 | 2.720.443 | 3.146.061 | 3.530.190 | 3.456.430 |
| L1-Tx1 | 1.332.071 | 707.047 | 492.758 | 1.328.940 | 247.067 |
| L2 | **24.295.832** | 18.778.236 | **18.931.729** | **15.671.956** | 10.981.470 |
| Penelope | 696.540 | 651.596 | 168.974 | 978.870 | 436.687 |
| Proto2 | 432.031 | 406.555 | 335.607 | 288.595 | 210.869 |
| R1 | 0 | 0 | 0 | 146.213 | 277.081 |
| R2-Hero | 258.797 | 37.021 | 152.400 | 101.778 | 204.860 |
| R2-NeSL | 139.492 | 0 | 191.055 | 106.574 | 62.871 |
| RTE | 1.012.896 | 0 | 0 | 0 | 0 |
| RTE-BovB | 3.915.100 | **11.058.368** | 4.053.280 | 5.607.772 | **8.522.404** |
| RTE-RTE | 0 | 0 | 0 | 169.573 | 0 |
| RTE-RTEX | 0 | 26.699 | 19.406 | 32.367 | 0 |
| RTE-X | 738.450 | 801.862 | 1.308.994 | 1.325.349 | 418.476 |
| Rex-Babar | **8.447.394** | 7.183.710 | **11.161.891** | **7.936.953** | 5.965.113 |
| Tad1 | 0 | 0 | 0 | 138.056 | 320.275 |
| LTR | 569.316 | 146.042 | 61.769 | 0 | 0 |
| Copia | 180.685 | 275.175 | 422.246 | 471.447 | 0 |
| DIRS | 766.684 | 743.991 | 1.555.092 | 1.105.144 | 825.655 |
| ERV1 | 750.875 | 1.123.315 | 980.225 | 1.469.763 | 126.774 |
| ERVK | 659.689 | 486.486 | 0 | 513.490 | 3.133.108 |
| Gypsy | 2.510.567 | 2.056.070 | 3.675.723 | 2.213.161 | 1.672.149 |
| Ngaro | 980.001 | 3.390.987 | 850.690 | 768.211 | 292.619 |
| Pao | 419.975 | 385.357 | 723.976 | 1.151.953 | 307.558 |
| RC | 0 | 0 | 0 | 0 | 0 |
| Helitron | **6.307.248** | 3.514.054 | **4.649.952** | **5.159.806** | 2.433.280 |
| Retroposon | 213.419 | 116.255 | 421.490 | 238.385 | 0 |
| SINE | 1.210.545 | 1.637.348 | 1.401.793 | 1.524.037 | 936.353 |
| 5S-Deu-L2 | 721.151 | 965.222 | 838.622 | 0 | 873.243 |
| ID | 0 | 0 | 19.012 | 29.728 | 116.970 |
| MIR | 1.185.268 | 1.020.008 | 1.163.355 | 2.103.257 | 382.739 |
| tRNA | 3.640.638 | 3.701.722 | 3.698.003 | 3.271.022 | 3.693.238 |
| tRNA-Core | 0 | 0 | 0 | 0 | 293.857 |
| tRNA-Core-L2 | 0 | 0 | 0 | 89.805 | 106.872 |
| tRNA-Deu-L2 | 0 | 0 | 0 | 42.616 | 0 |
| tRNA-L2 | 461.844 | 0 | 103.308 | 223.271 | 301.117 |
| tRNA-V | 0 | 0 | 492.471 | 0 | 0 |
| Unknown | 149.784.304 | 121.893.498 | 134.020.433 | 144.621.637 | 113.225.894 |
| **total interspersed** | **354.992.667** | **305.241.677** | **326.860.083** | **319.430.101** | **255.255.533** |
| Low_complexity | 2.964.797 | 2.285.494 | 2.321.578 | 2.478.451 | 2.097.985 |
| Satellite | 700.110 | 1.425.777 | 1.023.206 | 656.699 | 1.718.620 |
| Simple_repeat | 30.391.090 | 32.172.324 | 29.527.186 | 34.124.020 | 27.469.062 |
| rRNA | 358.199 | 351.617 | 313.179 | 42.963 | 708.978 |
| snRNA | 0 | 0 | 0 | 9.265 | 0 |
| **Total** | **389.406.863** | **341.476.889** | **360.045.232** | **356.741.499** | **287.250.178** |

## Notes

### Competing Interest Statement

The authors have declared no competing interest.

### Summary of Updates

Errors corrected for familly names of coauthors (and ORCID links added)

